# Open Source ImmGen: network perspective on metabolic diversity among mononuclear phagocytes

**DOI:** 10.1101/2020.07.15.204388

**Authors:** Anastasiia Gainullina, Li-Hao Huang, Helena Todorov, Kiwook Kim, Lim Sheau Yng, Andrew Kent, Baosen Jia, Kumba Seddu, Karen Krchma, Jun Wu, Karine Crozat, Elena Tomasello, Vipin Narang, Regine Dress, Peter See, Charlotte Scott, Sophie Gibbings, Geetika Bajpai, Jigar V. Desai, Barbara Maier, Sébastien This, Peter Wang, Stephanie Vargas Aguilar, Lucie Poupel, Sébastien Dussaud, Tyng-An Zhou, Veronique Angeli, J. Magarian Blander, Kyunghee Choi, Marc Dalod, Ivan Dzhagalov, Emmanuel L. Gautier, Claudia Jakubzick, Kory Lavine, Michail S. Lionakis, Helena Paidassi, Michael H. Sieweke, Florent Ginhoux, Martin Guilliams, Christophe Benoist, Miriam Merad, Gwendalyn J. Randolph, Alexey Sergushichev, Maxim N. Artyomov, ImmGen Consortium

## Abstract

We dissect metabolic variability of mononuclear phagocyte (MNP) subpopulations across different tissues through integrative analysis of three large scale datasets. Specifically, we introduce ImmGen MNP Open Source dataset that profiled 337 samples and extended previous ImmGen effort which included 202 samples of mononuclear phagocytes and their progenitors. Next, we analysed Tabula Muris Senis dataset to extract data for 51,364 myeloid cells from 18 tissues. Taken together, a compendium of data assembled in this work covers phagocytic populations found across 38 different tissues. To analyse common metabolic features, we developed novel network-based computational approach for unbiased identification of key metabolic subnetworks based on cellular transcriptional profiles in large-scale datasets. Using ImmGen MNP Open Source dataset as baseline, we define 9 metabolic subnetworks that encapsulate the metabolic differences within mononuclear phagocytes, and demonstrate that these features are robustly found across all three datasets, including lipid metabolism, cholesterol biosynthesis, glycolysis, and a set of fatty acid related metabolic pathways, as well as nucleotide and folate metabolism. We systematically define major features specific to macrophage and dendritic cell subpopulations. Among other things, we find that cholesterol synthesis appears particularly active within the migratory dendritic cells. We demonstrate that interference with this pathway through statins administration diminishes migratory capacity of the dendritic cells *in vivo*. This result demonstrates the power of our approach and highlights importance of metabolic diversity among mononuclear phagocytes.

## Introduction

The diversity of myeloid cells across different tissues is truly astonishing, both in function and in their developmental trajectory^1,2^. Additional dimension of this diversity is manifested by the metabolic characteristics of individual mononuclear phagocytes which can vary significantly based on the cell type and its location^3-5^. At present, direct metabolomics profiling of tissue residing subpopulations is not feasible, as the process of *ex vivo* sorting can be lengthy and cause significant metabolic perturbations^6,7^. However, RNA levels are significantly more stable to the sorting process and can serve as a reasonably reliable proxy to activities of metabolic pathways^8,9^. In this work we focus on understanding metabolic variability across phagocytic subpopulations through integrated examination of several large-scale datasets that transcriptionally profiled subsets of myeloid cells (**Fig. 1a-c**). Specifically, we have assembled compendium of three datasets, including the first public release of the new dataset generated by Mononuclear Phagocytes Open Source (MNP OS) ImmGen project^10^.

**Figure 1.**
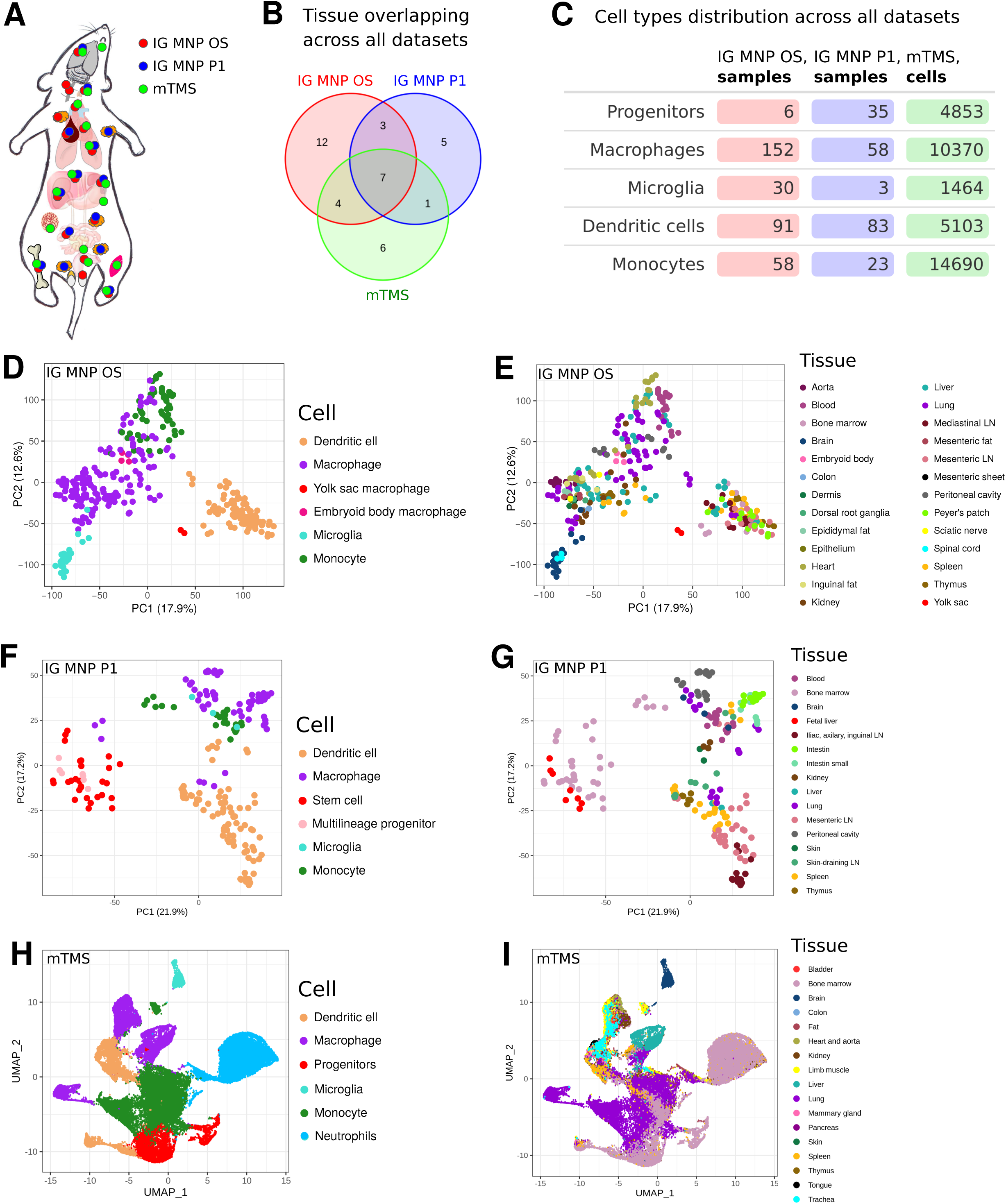
General overview of ImmGen Mononuclear Phagocytes Open Source (IG MNP OS), ImmGen Mononuclear Phagocytes Phase 1 (IG MNP P1) and myeloid Tabula Muris Senis (mTMS) datasets. **a**, Schematic representation of *Mus musculus* tissues where samples were derived from (marked with colored dots depending on the dataset). **b**, Number of tissues overlapping across all datasets. **c**, Cell types distribution across all datasets. Principal component analysis (PCA) based on 12,000 most expressed genes across all samples colored by the tissue of its origin (**e**,**g**) or cell type (**d**,**f**). UMAP representation of cells colored by the tissue of its origin (**i**) or its type (**h**). LN – lymph node.

ImmGen MNP OS dataset totals 337 samples and provides a unique source of information about individual cell subpopulations (**Fig. 1d, e**). It extends previous ImmGen effort that included 202 samples of various mononuclear phagocytes, also analysed in this study (**Fig. 1f, g**). In addition to increased number of mature cell populations from adult mice (monocytes, macrophages, and dendritic cells), the MNP OS dataset contains macrophages from the yolk sac (E10.5) and macrophages differentiated *in vitro* from embryonic stem cells (embryoid body derived macrophages, E6-E8). Furthermore, we leveraged recently released single-cell RNA-seq profiling of the multiple murine organs (Tabula Muris Senis^11^) and reanalysed those data by focusing only on the phagocytic populations, comprising 36,480 cells across 18 tissues (**Fig. 1h, i**). Taken together, a compendium of data assembled in this work covers multiple cell subpopulations found across 38 different tissues (**Fig. 1b**).

Using these transcriptional data, we sought to identify the major metabolic features characteristic of the different populations of phagocytic cells, and define how these features vary across the cell types and their locations. Such computational task has not been addressed previously for the datasets of such scale. Indeed, we previously described a computational approach, called GAM^12^, that uses metabolic networks as the backbone for analysis of transcriptional data and provides a verifiable and systematic description of the metabolic differences between *two conditions*^9^. However, datasets in question contain hundreds or even thousands of individual profiles, while GAM approach is designed to analyse comparison between two conditions. Therefore, in this work we have developed novel computational approach, GAM-clustering, which performs unbiased search of a collection of metabolic subnetworks that jointly define metabolic variability across large datasets (available on GitHub, see **Methods**). By doing so, GAM-clustering reveals metabolically similar subpopulations in a manner that does not require explicit annotation or pair-wise comparison of individual samples. Our analysis revealed major metabolic features associated with different cell subpopulations and highlighted a number of metabolic modules that are specific to individual cell types, tissues of residence, or developmental stages. As an example, GAM-clustering analysis revealed that cholesterol *de novo* synthesis pathway might play an important role in the context of migratory dendritic cells (DCs), which we validated using *in vivo* pharmacological inhibition of this pathway followed by tracking of DC migration. Consistent with the analysis, statins have demonstrated inhibitory effect on DC migratory ability, finding that has not been reported previously.

Taken together, our work provides both (1) unique data and analysis resource in terms of studying variability of mononuclear phagocytes, as well as (2) validated computational approach that can unbiasedly analyse both single-cell RNA-seq data as well as multi-sample bulk RNA-seq datasets in terms of key underlying metabolic features. Furthermore, we provide direct interactive access to the data for examination and visualization through both single-cell RNA-seq and bulk RNA-seq visualization servers including metabolic cluster annotations obtained in this work (https://artyomovlab.wustl.edu/immgen-met/).

## RESULTS

### Mononuclear Phagocytes Open Source (MNP OS) and Mononuclear Phagocytes from ImmGen Phase 1 (MNP P1) Datasets

As a part of the Open Source ImmGen Project, a total of 337 samples were collected and profiled through the collaborative effort of 16 laboratories (**Fig. 1d, e, Supplementary Fig. 1a, Supplementary Table 1**). Each laboratory sorted specific populations of mononuclear phagocytes from 26 distinct tissues, isolated RNA from these populations and submitted it for centralized deep RNA-sequencing and subsequent quantitation. Along with their samples of interest, each laboratory included RNA from locally sorted peritoneal macrophages as a common control for evaluation/correction of potential batch effects (**Methods**). Of note, 15 samples from MNP OS dataset were previously used in the study of sexual dimorphism of the immune system transcriptome^13^, while complete dataset is analysed in this work for the first time.

Overall, the transcriptional data demonstrated high concordance between different collection sites, and were merged into a final transcriptional master table (**Supplementary Fig. 1b, c, Supplementary Table 2, Methods**). Previously established markers of individual myeloid subpopulations^14–18^ matched well with the sorted populations (**Supplementary Fig. 2**), indicating overall consistency of the dataset across different research groups. As individual principal component analysis (PCA) plots show (**Fig. 1d**), samples have clustered in accord with their broad annotation as macrophages, DCs, monocytes or microglia. Generally, subpopulation-specific effects were stronger than tissue specific differences within individual subpopulations as evident by comparing **Figures 1d** and **1e**. To estimate the degree of metabolic variability in the data, we examined the enrichment of annotated metabolic pathways in this dataset, revealing coherent transcriptional patterns across individual subpopulations (**Supplementary Fig. 3a**). This indicated that systematic evaluation of the metabolic subnetworks within the data is warranted.

**Figure 2.**
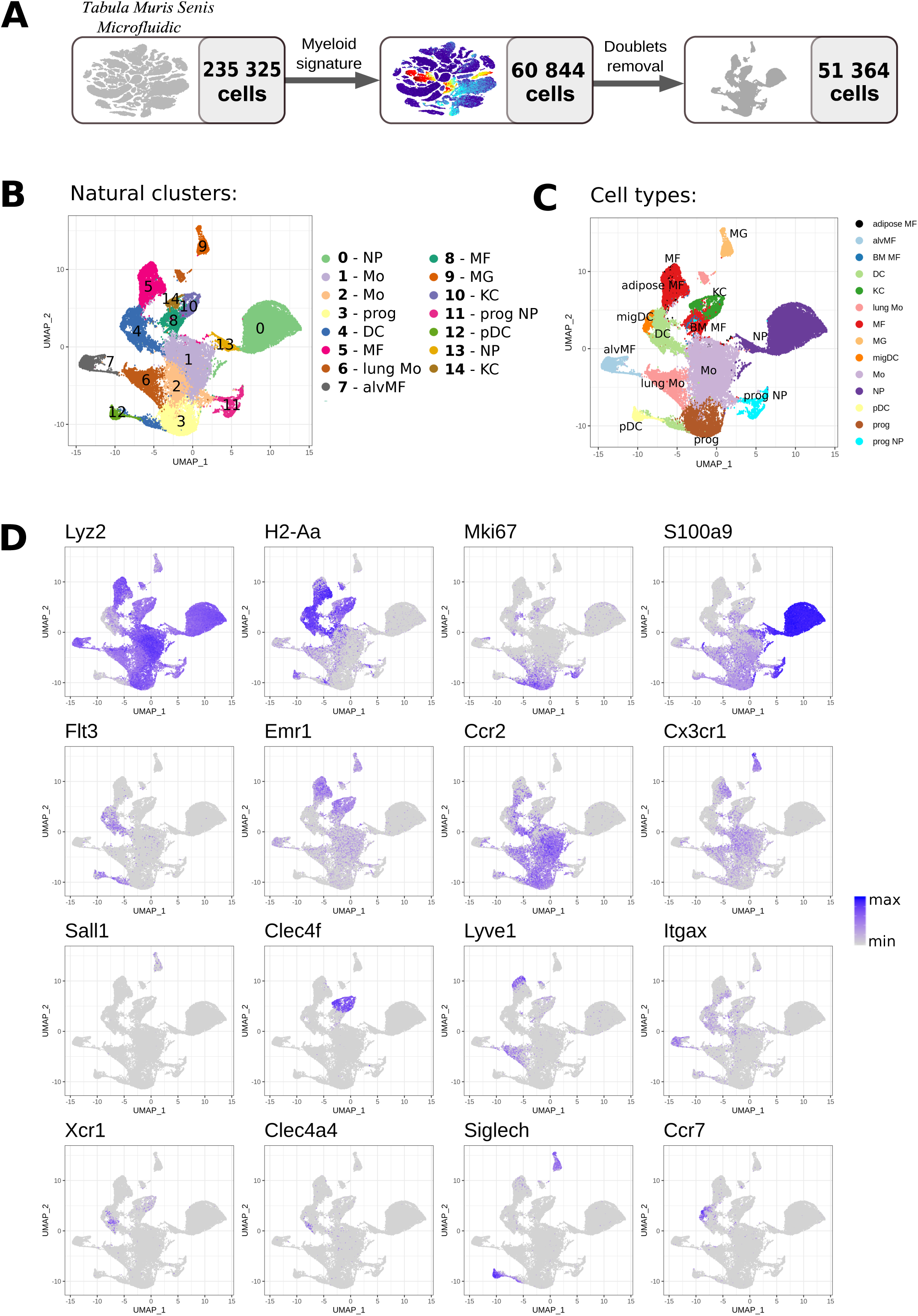
Tabuls Muris Senis single cell RNAseq dataset. **a**, Dataset preprocessing resulting in myeloid subset derivation. UMAP plot with natural clusters (**b**) and cell types (**c**) identified based on cell specific markers (**d**). NP – neutrophil, Mo – monocyte, prog – progenitor, DC – dendritic cell, MF – macrophage, alv MF – alveolar macrophage, MG – microglia, KC – Kupffer cell, pDC – plasmacytoid dendritic cell, migDC – migratory dendritic cell.

**Figure 3.**
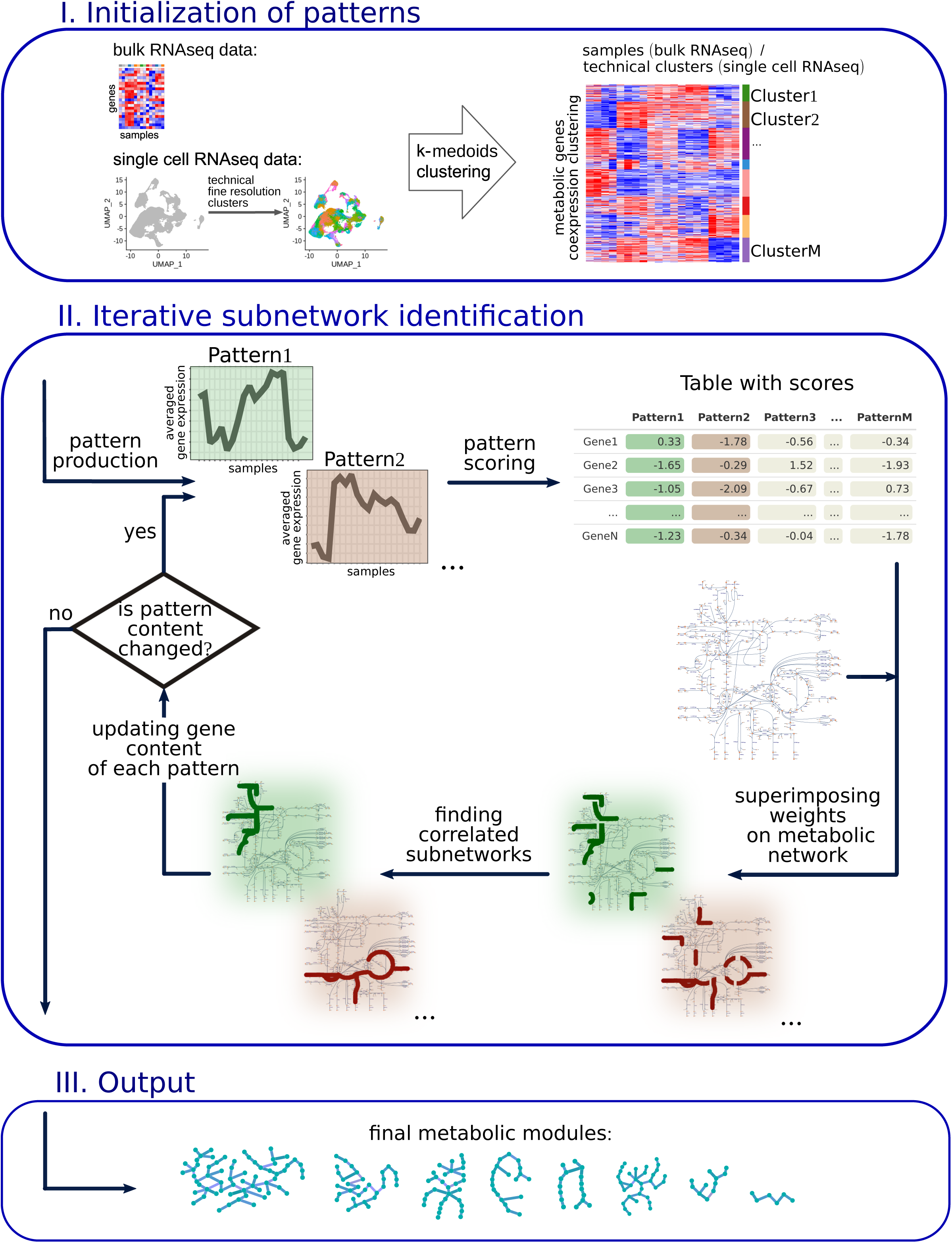
Scheme of analysis approach for multisample metabolic network clustering (GAM-clustering). The dataset’s metabolic genes are initially clustered based on a k-medoids algorithm. Averaged gene expression of the obtained clusters is further considered as patterns. For each gene, a score is calculated on the basis of its correlation with each pattern. These scores are superimposed on the KEGG metabolic network. Based on these scores, the most weighted connected subnetwork is found for each parent. After the refinement procedure, metabolic modules as a final version of subnetworks are obtained.

Initial ImmGen Phase 1 data published previously^5^ include 202 samples of mononuclear phagocytes with higher contribution of progenitor populations, and smaller number of microglial samples (**Fig. 1f**) overall spanning 16 tissues (**Fig. 1g**) – we will refer to this subset as ImmGen MNP P1 from here onward. Similar to MNP OS dataset, enrichment in metabolic pathways across subpopulations in MNP P1 data has demonstrated coordinate variations across the samples (**Supplementary Fig. 3b**).

### Single-cell Myeloid Tabula Muris Senis (mTMS) dataset

Tabula Muris consortium has performed single-cell RNA-sequencing for a large number of tissues without explicit sorting into individual cell populations^11^. These data include myeloid cells localized in the corresponding tissues which can be computationally separated based on the expression of common myeloid signatures. Using latest public dataset, Tabula Muris Senis, we have analysed the data for 235,325 cells to identify 51,364 myeloid cells (mononuclear phagocytes and neutrophils) that expressed key myeloid markers (Lyz2, H2-Aa, Mki67, S100a9, Flt3, Emr1, Ccr2, Cx3cr1, Sall1, Clec4f, Lyve1, Itgax, Xcr1, Clec4a4, Siglech, Ccr7, **Fig. 2a**,**d, Methods**). These cells comprised dedicated dataset, further referred to as myeloid Tabula Muris Senis dataset (mTMS). While single-cell RNA-seq data inevitably detect smaller number of genes per cell compared to bulk RNA-sequencing (**Supplementary Fig. 4**), the depth of the mTMS dataset was sufficient to resolve classical cell populations. Specifically, unbiased clustering revealed 15 subpopulations within mTMS dataset (**Fig. 2b**), which could be readily identified as well described populations of plasmacytoid DCs, monocytes, Kupffer cells, microglia etc (**Fig. 2c**) based on previously described cell-specific markers (**Fig. 2d**). To our knowledge, we provide the first large-scale curated annotation for the myeloid cells within Tabula Muris Senis data. Corresponding annotations are available for hands-on exploration in the interactive single-cell browser (http://artyomovlab.wustl.edu/immgen-met/, see Tabula Muris Senis).

**Figure 4.**
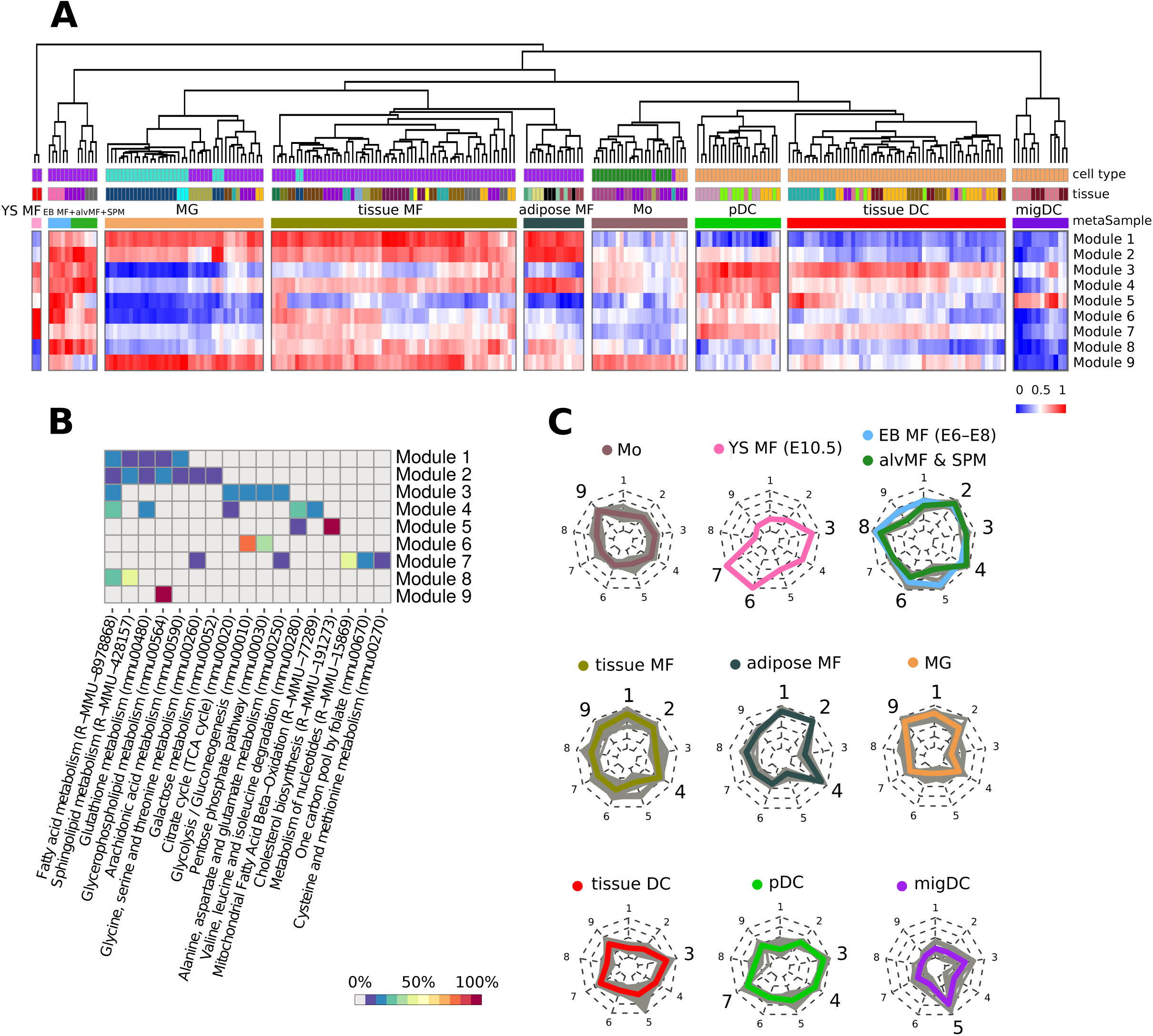
Metabolic modules as a result of multisample metabolic network clustering of all myeloid cells but not inflammatory conditions from ImmGen MNP OS dataset. **a**, Heatmap representing samples hierarchically clustered based on averaged gene expression of each of obtained module (from lowest as blue to highest as red). Euclidean distance is used as a clustering metric. YS MF – yolk sac macrophage, EB MF – embryoid body macrophage, alvMF – alveolar macrophage, SPM – small peritoneal macrophage, MG – microglia, MF – macrophage, Mo – monocyte, DC – dendritic cell, pDC – plasmacytoid DC, migDC – migratory DC. **b**, Annotation of the obtained modules based on gene enrichment in KEGG and Reactome canonical pathways. Enrichment value is calculated as a percentage of module genes contained in a particular pathway. **c**, Radar chart representation of metabolic modules within each metasample. Each individual sample is shown as a grey line while mean of all samples inside one metasample is shown as a colored line. Nine radii of radar chart are devoted to the corresponding metabolic modules: 1,2 – Lipid metabolism, 3 – FAS pathway, 4 – mtFASII pathway, 5 – Cholesterol synthesis, 6 – Glycolysis, 7 – Folate, serine & nucleotide metabolism, 8 – FAO & sphingolipid *de novo* synthesis, 9 – Glycerophospholipid metabolism. Metasamples of EB MFs + alvMFs and alvMFs + SPMs cells are shown at one chart as they are extremely close in their metabolic characteristics.

### GAM-clustering: identification of metabolic subnetworks in datasets with multiple conditions

Previously, we have shown that metabolic remodelling between two conditions can be analysed using network-based analysis of their transcriptional profiles^9,12^. Specifically, the GAM (‘Genes and Metabolites’) algorithm searches for optimal subnetworks within a global metabolic network by weighing individual enzymes in accord with differential expression of their genes and then solving the generalized maximum-weight connected subgraph (GMWCS) problem^12,19^. While this approach cannot be directly translated to multi-sample/single-cell datasets such as ImmGen or Tabula Muris Senis data, we were able to reformulate weighting scheme in a manner that allows GMWCS subnetwork search without explicit annotation of individual samples or conditions. Here, we describe novel algorithm called GAM-clustering that allows the user to obtain metabolic subnetworks enriched within the transcriptional data that include many samples across multiple conditions.

In brief, GAM-clustering searches for connected metabolic subnetworks that have most correlated expressions of individual enzymes, resulting in a collection of subnetworks that follow distinct transcriptional profiles. To achieve that, we first initialize a pattern-generating profiles by clustering all metabolic genes based on their co-expression patterns (**Fig. 3**, see **Methods** and **Supplementary Material** for details). For initialization of the multisample bulk RNA-seq data we use k-medoids clustering with k=32 (see **Supplementary Material** and **Supplementary Fig. 5a**,**b** for parameters sensitivity), any other gene expression clustering approach can be used in this step since downstream steps include significant re-grouping and merging of individual clusters. For initialization of single-cell RNA-seq data, we first cluster the cells in the dataset to multiple clusters (∼100) which provides sufficient balance between fine resolution of the data and minimal coarse-graining needed to avoid drop-out artefacts (see **Methods** and **Supplementary Material** for details). Then, genes are clustered using the same procedure as for multisample bulk datasets.

**Figure 5.**
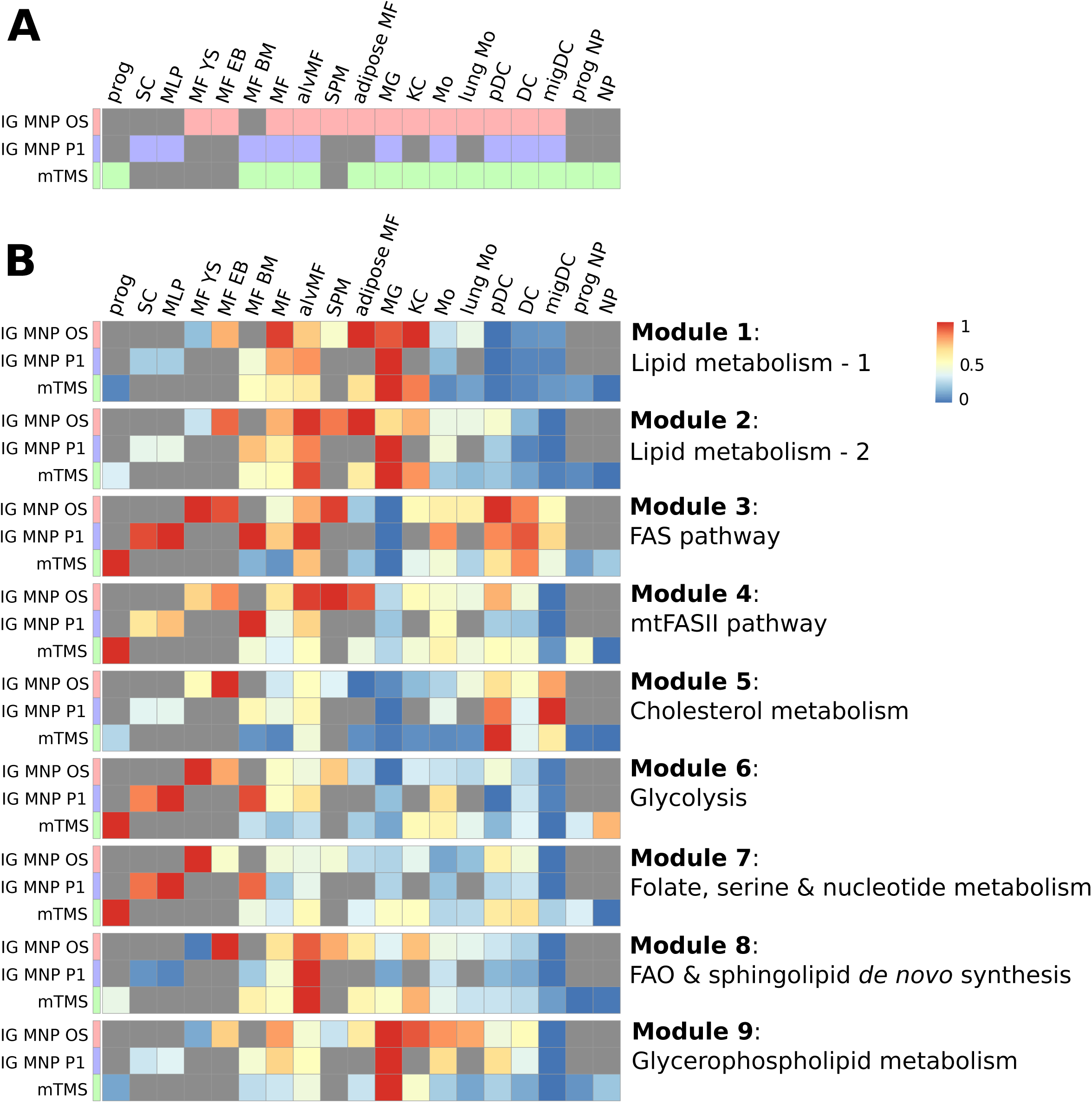
Cell types shared between ImmGen Mononuclear Phagocytes Open Source (IG MNP OS), ImmGen Mononuclear Phagocytes Phase 1 (IG MNP P1) and myeloid Tabula Muris Senis (mTMS) datasets have similar patters of metabolic modules signatures. **a**, Population memberships across the datasets: prog – progenitor, SC – stem cell, MLP – multilineage progenitor, MF YS – yolk sac macrophage, MF EB – embryoid body macrophage, MF – macrophage, alvMF – alveolar macrophage, SPM – small peritoneal macrophage, MG – microglia, KC – Kupffer cell, Mo – monocyte, pDC – plasmacytoid dendritic cell, DC – dendritic cell, migDC – migratory dendritic cell, NP – neutrophil (**Supplementary Table 4**). **b**, Enrichment of individual metabolic modules across all datasets obtained during GAM-clustering analysis of IG MNP OS dataset.

Next, each enzyme is weighed with respect to the similarity of its transcript’s profile to the average cluster/pattern profile, resulting in multiple weights per gene that are specific for each pattern (**Fig. 3**). For any given pattern, weights of individual enzymes serve as input to the GMWCS solver, resulting in the individual subnetworks that are associated with each pattern. Individual subnetworks are then refined in an iterative procedure of updating the gene content for each pattern (see **Supplementary Material**). The final output presents a set of specific subnetworks that reflect metabolic variability within a given transcriptional dataset (**Fig. 3**).

### Major metabolic modules within mononuclear phagocyte subpopulations

The GAM-clustering algorithm was applied to data from all baseline (non-infected) samples and yielded nine distinct metabolic modules (**Fig. 4a, Supplementary Table 3**). Hierarchical clustering of samples based on the Euclidean distance metric in space of these nine metabolic modules showed that they can be broadly separated based on the cell types: yolk sac macrophages, dendritic cells, monocytes, and macrophages from adult organism. Broadly defined mononuclear cell types are further split into several smaller metasamples: dendritic cells subdivided into plasmacytoid dendritic cells (pDCs), tissue specific dendritic cells, and migratory dendritic cells (migDCs), and macrophages subdivided into microglia, adipose tissue macrophages, and a large metasample of tissue residing macrophages, as well as an additional metasample composed of embryoid body, alveolar, and small peritoneal macrophages (SPMs) that clustered distinctly from other macrophage subpopulations (**Fig. 4a**).

While obtained metabolic modules/subnetworks provide a more accurate description of metabolic diversity compared to canonically annotated pathways, the latter can be useful for coarse-grained understanding of functionalities associated with each subnetwork (**Fig. 4b**). Indeed, pathway enrichment analysis along with subnetworks gene content analysis indicate that modules 1, 2, 8 and 9 represent various aspects of lipid metabolism, and modules 3, 4 – two types of fatty acid synthesis pathways. Finally, distinct modules represent cholesterol synthesis metabolism (module 5), glycolysis (module 6) and nucleotide/folate metabolism associated subnetworks (module 7).

The underlying metabolic phenotypes for each metasample can be represented using radar chart diagrams (**Fig. 4c**): each metasample is defined by a specific combination of metabolic features that provides unique insights into metabolic wiring within those populations. Here, the names to metasamples are given based on the most common sample type inside the cluster. An alternative view of the samples in the space of metabolic modules can be obtained using PCA that is built based on only 9 metabolic modules, which shows distinct separation of individual metasamples (**Supplementary Fig. 6a**). Consistently, when overlaid with the PCA representation from **Figure 1**, individual metabolic modules formed coherent patterns indicating the groups of metabolically similar samples (**Supplementary Fig. 6b**). Altogether, the metabolic modules/subnetworks and corresponding metasamples encapsulate metabolic variability across both cell types and their tissues of residency. We next turn to examining robustness of the obtained subnetworks across three considered datasets.

**Figure 6.**
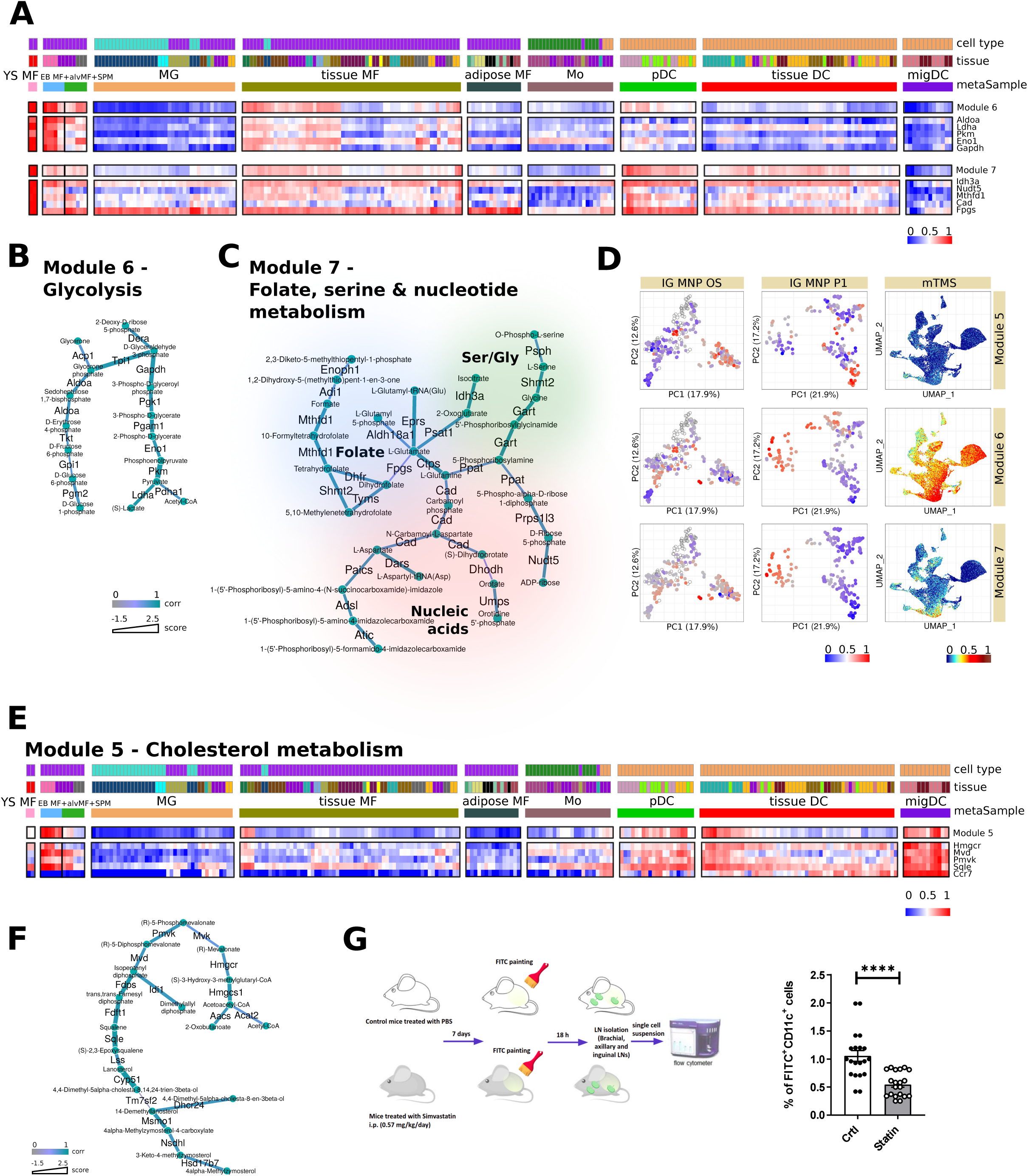
Subnetworks associated with early developmental stages and dendritic cells. Heatmaps of module patterns along with the expression of some of its genes or genes related to the same biological subject (from lowest as blue to highest as red). YS MF – yolk sac macrophage, EB MF – embryoid body macrophage, alvMF – alveolar macrophage, SPM – small peritoneal macrophage, MG – microglia, MF – macrophage, Mo – monocyte, DC – dendritic cell, pDC – plasmacytoid DC, migDC – migratory DC (**a**,**e**). Metabolic modules *per se* where edges of modules are attributed with color according to correlation of its enzyme’s gene expression to this particular module pattern and thickness according to its score (**b**,**c**,**f**). **d**, Enrichment of modules genes expression (from lowest as blue to highest as red, transparent dots correspond to treated samples) across all three analysed datasets: ImmGen Mononuclear Phagocytes Open Source (IG MNP OS), ImmGen Mononuclear Phagocytes Phase 1 (IG MNP P1) and myeloid Tabula Muris Senis (mTMS) datasets. **g**, Dendritic cell migrations experiment scheme and results.

### Three independent large-scale datasets show consistent metabolic features

We next considered if metabolic subnetworks derived from ImmGen MNP OS data can be seen in the other two large scale datasets considered in this work – ImmGen MNP P1 and mTMS datasets. While overlap in profiled tissues is considerate (**Fig. 1b**), 3 datasets are not identical in terms of populations profiled. We, therefore, grouped the samples into 19 general classes and compared the datasets by looking at the metabolic enrichments across these classes (**Fig. 5a, Supplementary Table 4**). To examine robustness of metabolic signatures, we computed enrichments of individual metabolic modules from **Figure 4a** in each of the 19 representative classes of ImmGen MNP OS, ImmGen MNP P1 and mTMS. Indeed, all datasets modules demonstrated extremely similar enrichment profiles (**Fig. 5b**): for instance, Module 1 was enriched in microglia, adipose tissue macrophages and Kuppfer cells, module 8 enriched in alveolar macrophages, module 5 – in pDCs and migratory DCs across all datasets, etc.

Importantly, independent application of the GAM-clustering algorithm to each of the datasets also revealed very high degree of similarity in obtained modules, highlighting reproducible and robust nature of the derived metabolic subnetworks (**Supplementary Fig. 7, Supplementary Table 3, Supplementary Material**).

**Figure 7.**
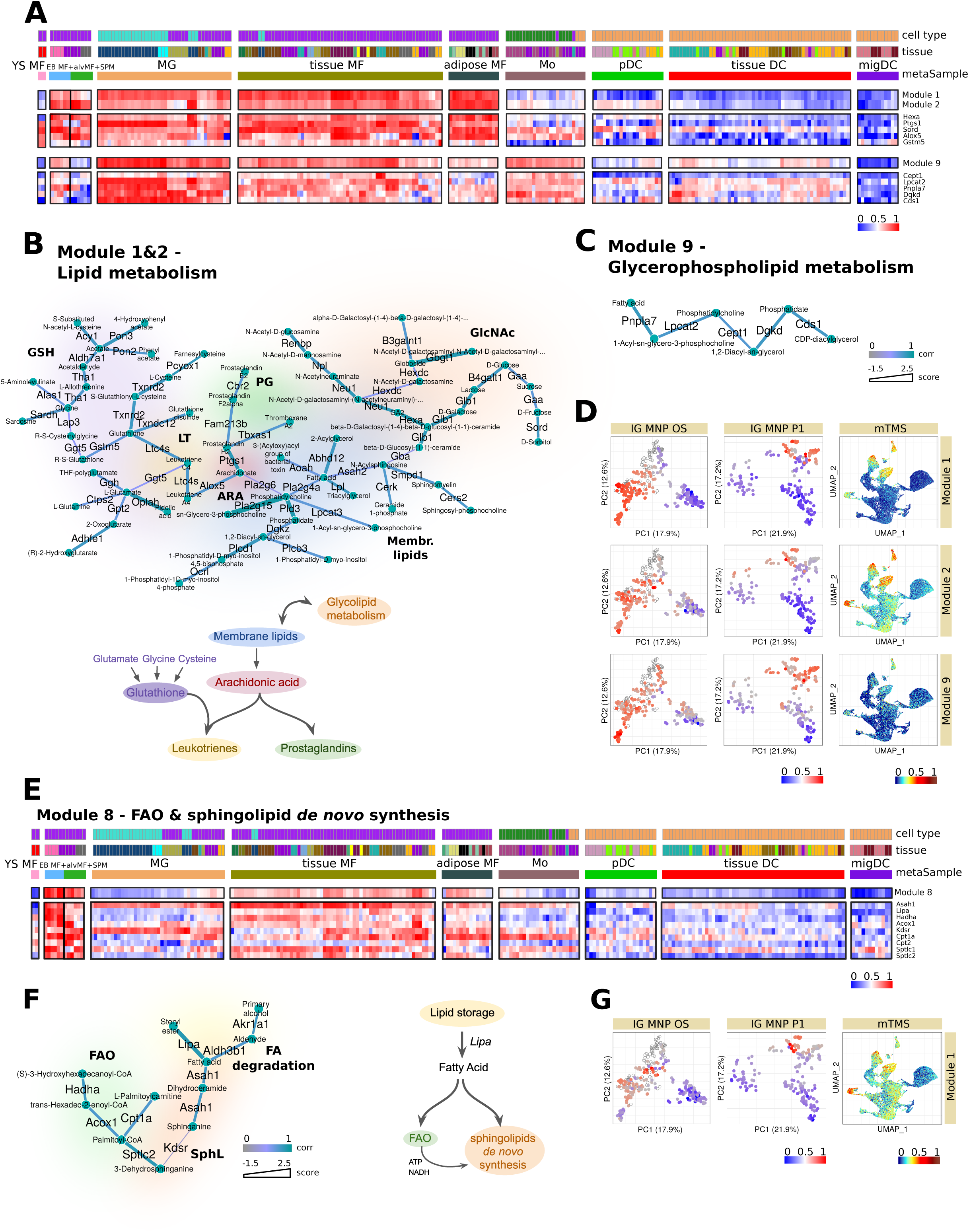
Subnetworks associated with fatty acid synthesis and degradation. Heatmaps of module patterns along with the expression of some of its genes (from lowest as blue to highest as red). YS MF – yolk sac macrophage, EB MF – embryoid body macrophage, alvMF – alveolar macrophage, SPM – small peritoneal macrophage, MG – microglia, MF – macrophage, Mo – monocyte, DC – dendritic cell, pDC – plasmacytoid DC, migDC – migratory DC (**a**,**e**). Metabolic modules *per se* and corresponding schematic diagrams. Edges of modules are attributed with color according to correlation of its enzyme’s gene expression to this particular module pattern and with thickness according to its score. (**b**,**c**,**f**). Enrichment of modules genes expression (from lowest as blue to highest as red, transparent dots correspond to treated samples) across all three analysed datasets: ImmGen Mononuclear Phagocytes Open Source (IG MNP OS), ImmGen Mononuclear Phagocytes Phase 1 (IG MNP P1) and myeloid Tabula Muris Senis (mTMS) datasets (**d**,**g**).

Next we examined individual subnetworks from the perspective of metabolic reactions covered and describe both published evidence of the corresponding metabolic activities as well as validation data obtained in this project.

### Subnetworks associated with early developmental stages

Module 6 (**Fig. 6a**,**b**,**d**) is one of the modules most distinctly associated with yolk sac, embryoid body, alveolar, and small peritoneal macrophages. This module, though unbiasedly derived by our network analysis, closely matches the canonical glycolysis pathway (**Fig. 6b**), indicating strong transcriptional co-regulation of these genes across the collected samples. Enrichment of the glycolysis module in developmental cell types is consistent with previously published data highlighting the importance of glycolysis for stem-like and progenitor populations^20–23^. This is also consistent with the ImmGen MNP P1 and mTMS data (**Fig. 6d**), where this module is also most enriched in progenitor populations. Interestingly, mTMS single-cell RNA-seq data also demonstrate that this module is enriched in neutrophils, in accord with described high glycolytic rate in these cells^24^.

Module 7 (**Fig. 6a**,**c**,**d**) represents another set of metabolic activities including folate and serine metabolism as well as the nucleotide biosynthesis pathway typically associated with the progenitor populations^25–27^. In addition to the yolk sac macrophages, this module is also enriched in some tissue residing dendritic cells and pDCs (but not in migDCs). Indeed, the importance of some of these pathways (e.g. folate metabolism) has been demonstrated in dendritic cell functions such as antigen presentation^28^.

### Cholesterol synthesis pathway is enriched in and functionally important for migratory DCs

Module 5 almost exclusively consists of enzymes from the cholesterol metabolism/mevalonate synthesis pathway (**Fig. 6d**,**e**,**f**), and is enriched in embryoid body macrophages and some dendritic cells. Specifically, cholesterol synthesis appears to play a major role in migDCs, while it is less prominent in pDCs and conventional tissue residing dendritic cells. Additionally, with respect to potential tissue-specific imprinting, it is worth noting that a small subset of tissue residing macrophages, comprised of epithelial and dermal macrophages, is enriched in genes of the mevalonate/cholesterol synthesis pathway (**Fig. 6e**). Enrichment of cholesterol metabolism in migratory dendritic cells was consistent with mechanistic data by Hauser and colleagues who showed that cellular cholesterol levels are directly linked to the ability of dendritic cells to oligomerize Ccr7 (key marker of migDCs) and acquire a migratory phenotype^29^. Given the results of our analysis and these published mechanistic connections, we evaluated mobilization of DCs to lymph node following epicutaneous application of fluorescein isothiocyanate (FITC) in either control animals or animals treated i.p. with low-dose Simvastatin (0.57 mg/kg/day) for seven days (**Fig. 6g**). Draining lymph nodes collected 18 h after FITC application demonstrated significantly fewer migrated FITC^+^CD11c^+^ dendritic cells in the animals treated with Simvastatin, illustrating that *in vivo* interference with cholesterol synthesis reduces dendritic cell migration to the lymph node, fitting with the prominent expression of cholesterol synthesis genes in DCs. This results illustrates general validity of our analysis and highlights novel features of the systemic metabolic perturbations, such as statin treatments, that we previously not recognized.

### Subnetworks associated with lipid metabolism

Modules 1 and 2 cover various aspects of lipid metabolism and are strongly specific to macrophages relative to monocytes and dendritic cells (**Fig. 7a**). Due to general similarity of their patterns, we merged the subnetworks for modules 1 and 2 in order to make the interpretation easier (**Fig. 7b, Supplementary Fig. 8a**,**b**). The resulting subnetwork is centred around phospholipid and arachidonic acid metabolism, and includes parts of the glutathione and cysteine/glutamate/glycine metabolism pathways, as well as the N-acetylglucosamine pathway. Indeed, arachidonic acid metabolism has been shown to play major roles in macrophages^30,31^. Its metabolic flow is associated with utilization of phospholipids to produce two major classes of the arachidonic acid derivatives: leukotrienes and prostaglandins. Unlike prostaglandins, leukotriene production (C4 and downstream) requires glutathione as an intermediate metabolite, thus involving the glycine, cysteine and glutamate pathways^32^. Furthermore, our analysis picked up a distinct subnetwork of co-expressed genes from the glycerophospholipid pathway (Module 9, see **Fig. 7a**,**c**,**d**) that was particularly highly expressed in the microglial populations (**Fig. 7a**). This module included enzymes such as Dgkd and Lpcat2, suggesting that their role in microglia might be of particular interest^33,34^. As **Fig. 7d** shows, these observations were common across all three datasets.

### Subnetworks associated with fatty acid synthesis and degradation

Our analysis identified three distinct subnetworks associated with modulation of fatty acids in terms of both their synthesis (modules 3 and 4) and fatty acid oxidation (module 8).

The structure of Module 3 (**Supplementary Fig. 8a**,**b**,**d**) reflects energetic demands of the fatty acid synthesis and includes portions of pentose phosphate pathway and TCA cycle, where citrate synthase (Cs) is one of the most pattern-specific genes within this subnetwork. Overall, module 3 is highly enriched in dendritic cell populations, but not in macrophage/monocyte samples, underscoring another facet of metabolic divergence between these cell types. The functional importance of this module for dendritic cells is evident from the fact that a blockade of Fasn-mediated fatty acid synthesis markedly and selectively decreases dendropoiesis both in mice and in humans^35,36^.

Interestingly, the pattern of Module 8 (**Fig. 7e-g**) was directly opposite to Module 3, and was strongly enriched among various tissue macrophages, particularly in alveolar macrophages. Metabolic flow encompassed by this network includes enzymes such as Lipa (LAL), which is responsible for lysosomal lipolysis and initial breakdown of intracellular lipid storage. This breakdown is followed by mitochondrial import of cytosolic fatty acids via carnitine transport shuttle (Cpt1a) and their subsequent breakdown via classical FAO steps (Acox1, Hadha, etc)^37,38^ (**Fig. 7e**). The Lipa expression pattern is one of the most specific for module 8, indicating its potential importance for macrophages. Indeed, there are studies highlighting the importance of Lipa for macrophage function, especially in the context of anti-inflammatory polarization^39^. Furthermore, Lipa is also likely to be important for human macrophages, as mutations in the LIPA gene of patients with cholesteryl ester storage disease (CESD) cause aberrant cholesterol accumulation in tissue macrophages^40,41^. Enrichment of fatty acid oxidation-related Module 8 in alveolar macrophages is particularly interesting as it is distinctly reproduced in ImmGen MNP P1 and mTMS data. Importance of this pathway in lungs is intriguing and warrants further detailed investigations.

## DISCUSSION

Here we introduced unique dataset covering multiple subpopulations of dendritic cells, monocytes and macrophages from diverse tissues – result of ImmGen MNP Open Source profiling effort. We focused on understanding potential metabolic variability among collected myeloid cell subpopulations and co-analysed it in the context of two other large-scale profiling efforts – ImmGen Phase 1 and Tabula Muris Senis. Using new algorithmic approach (GAM-clustering), we have defined 9 major metabolic subnetworks that encapsulate the major metabolic differences that were highly reproducible across three studied datasets. Our analysis demonstrated that specific metabolic features could be attributed to cell populations as well as specific tissue of residence for distinct populations (e.g. adipose tissue macrophages).

Our analysis suggested that major metabolic differences between baseline (unactivated) macrophages and dendritic cells are (1) levels of Fasn-mediated fatty acid synthesis enriched in dendritic cells’ transcriptional profiles, and (2) regulation of arachidonic acid metabolism, which is enriched in macrophages. Among various tissue residing cell types, it was apparent that microglia and CNS macrophages have a very distinct phenotype relative to other populations: based on their transcriptional profile they appear more metabolically quiescent, yet a particular lipid-associated module (module 9) was enriched in these cells, with key genes being Lpcat2, Dgkd, Csd1 that are involved in phospholipid metabolism and the generation of bioactive lipids from phospholipid precursors.

Indeed, distinct patterns in lipid metabolism, including pathways related to cholesterol, were also apparent in dendritic cells versus macrophages. Macrophages’ capacity to handle cholesterol and store it in esterified form to generate so-called macrophage foam cells is a well-established theme in cardiovascular research and inflammatory disease^42,43^. Our data reveal that expression of Lipa, an enzyme involved in breaking down cholesterol esters in the lysosome and whose mutation is associated with lysosomal storage diseases, is a widespread characteristic of tissue macrophages but not dendritic cells. On the contrary, we observed that pathways active in cholesterol synthesis are very low in all tissue macrophages but elevated in monocytes and dendritic cells, especially migratory dendritic cells.

Thus, it appears as though macrophages are oriented toward handling exogenously derived cholesterol, such as that which may be derived from engulfment of large amounts of phagocytic cargo, whereas dendritic cells are oppositely programmed to synthesize their own cholesterol and associated intermediates. As tissue macrophages are especially incapable of migrating to distal sites like lymph nodes, a major functional distinction from dendritic cells, we have validated importance of the cholesterol synthesis pathway for the migratory phenotype *in vivo* by using pharmacological interventions with Simvastatin.

Altogether, our analysis underscores metabolic variability across cell types and tissues and highlights the need to understand metabolic wiring, not only in terms of cellular metabolism, but also at the level of whole-body communication networks (see e.g. Castillo-Armengol and colleagues^44^, Droujinine and Perrimon^45^). Furthermore, since direct metabolic profiling is not feasible or sufficiently accurate at the moment, the development of *ex vivo* metabolomics profiling technologies^46–48^ suggests that direct insight into metabolism of various myeloid subpopulations through *in vivo* metabolomics techniques will be possible in future.

## METHODS

### RNA-sequencing

Bulk RNA-sequencing data were collected from 16 labs. All of the mice used in this study were handled in accordance with IACUC-approved protocols. Each lab, in addition to their own samples, sorted a standard peritoneal cavity macrophage population (CD115^+^B220^-^ F4/80^hi^MHCII^-^) for comparability between all labs. Samples were profiled using ImmGen’s ultra low input (ULI) sequencing pipeline, in batches of 90-96 samples. All samples were sequenced in two separate NextSeq500 runs and combined for increased depth (expect 8-12 10^6^ reads per sample). Following sequencing, raw reads were aligned with STAR to the mouse genome assembly mm10, and assigned to specific genes using the GENCODE vM12 annotation. Aligned reads were quantified using featureCounts. Samples that did not pass the QC threshold for read counts (<2 million reads) were dropped for further analysis. Pearson correlation was calculated between biological replicates to exclude samples that did not pass a threshold of 0.9 correlation coefficient. For the cell populations with three biological replicates, of which one did not agree with the other two, the suspect one was removed from the data set. In case cell populations had only two replicates, both were removed. Samples with Jchain>1,000 and Ighm>10,000 were set asides as well as samples with high B cell, erythrocytes and fibroblasts transcripts. Peritoneal cavity samples were downsampled to keep consistency across samples number in all tissues.

### RNA-sequencing data processing

All gene counts were imported into the R/Bioconductor package EdgeR and TMM normalization size factors were calculated to adjust for differences in library size across all samples. Feature not expressed in at least three samples above one count-per-million were excluded from further analysis and TMM size factors were recalculated to create effective TMM size factors. The effective TMM size factors and the matrix of counts were then imported into the R/Bioconductor package Limma and weighted likelihoods based on the observed mean-variance relationship of every gene and sample were then calculated for all samples. Performance of the samples was assessed with a Pearson correlation matrix and multidimensional scaling plots.

### Single-cell RNA-seq data processing

Filtered h5ad file for Droplet subset was downloaded from the official Tabula Muris Senis repository (https://figshare.com/projects/Tabula_Muris_Senis/64982). The data were processed by the standard Seurat pipeline and resulted in 235,325 cells organised in distinct clusters detectable on TSNE/UMAP plots. Next, cells annotated with names corresponding to myeloid populations were picked out. A differential gene expression analysis between these cells and all others was performed. Top 250 of these differentially expressed genes were used as a “myeloid signature genes” (**Supplementary Table 5**) to identify clusters that most express them and thus correspond to myeloid cells. Cell content of these clusters was used to create a new subset of 60,844 cells. Obtained dataset was analysed by non-myeloid marker genes to detect and remove cell doublets with T-cells, B-cells, NK-cells and fibroblasts (Cd3d, Cd3e, Cd3g, Cd4, Cd8a, Cd19, Cd79a, Tnfrsf17, Cd22, Nkg7, Gnly, Col6a1, Col6a2, Col6a3). Finally, dataset of 51,364 cells was obtained and used in the further GAM-clustering analysis.

### GAM-clustering

The algorithm for multisample metabolic network clustering (hereinafter referred to as GAM-clustering, see **Supplementary Material** and **Supplementary Fig. 5** for details) identifies modules describing dynamic regulation of metabolism and is based on the previously developed GAM method^12^. GAM-clustering extends the GAM method by setting the task to find not one but several metabolic modules (connected subnetworks of metabolic network) with the condition that each of these modules should contain as many metabolic genes with high pairwise correlation of their expression as possible.

The initial approximation of the final set of modules is carried out by k-medoids clustering of a gene expression matrix for all metabolic genes of a dataset with some arbitrary k (here we used k=32). Each cluster forms a corresponding expression pattern which can be determined as averaged value of its z-normalized gene expression values. The metabolic network used for further analysis is presented as a graph where vertices are metabolites and edges are KEGG database reactions which are mapped with catalysing them enzymes and corresponding genes. For each particular pattern edges of this graph are scored (weighted) based on their gene expression similarity with this pattern and dissimilarity with other patterns.

For each case of weighted graph a connected subgraph of maximal weight is found by a signal GMWCS (generalized maximum weight connected subgraph) solver^49^ (https://github.com/ctlab/sgmwcs-solver) and is called a metabolic module. This solver uses the IBM ILOG CPLEX library, which efficiently performs many iterations of this method in a reasonable amount of time. Then, each pattern is updated by replacing it with an averaged gene expression of the module’s edges with a positive score. If the pattern is changed, a new score set is calculated and a new iteration is performed. Before moving to the next iteration, small graphs are eliminated from further analysis so that there are no graphs with less than five edges and diameter less than four in the output solution. The algorithm continues until the pattern content stops changing.

GAM-clustering method is applicable not to bulk RNA-seq data only but to single-cell RNA-seq data as well. Single-cell data need an additional step of preprocessing implying transformation of individual cells into technical samples. This is performed based on averaging gene expressions of individual cells inside high resolution clusters. In case of single-cell RNAseq data, among final metabolic modules might occur ones that do not cover all biological replicas of cell types they are specific for. These modules are eliminated from the final result.

Thus, the final metabolic modules are subnetworks of the overall metabolic network that contain a set of closely located genes with high correlation of their expression profile across all samples.

GAM-clustering method is available at https://github.com/artyomovlab/ImmGenOpenSource.

### DC migration

Epicutaneous application of Fluorescein isothiocyanate (FITC) to study DC migration was performed on three areas of each side of the mouse back skin as described previously ^50^. Both females and males were studied. Briefly, FITC (8 mg/ml) was dissolved in acetone and dibutyl phthalate (Sigma-Aldrich, F7250) and applied in 25-μl aliquots to each site. Recovered LNs, 18l aliquots to each site. Recovered LNs, 18 h later, were teased and digested in 2.68 mg/ml collagenase D (Roche) for 25 min at 37°C. Then, 100 μl aliquots to each site. Recovered LNs, 18l 100 mM EDTA was added for 5 min, and cells were passed through a 100-μl aliquots to each site. Recovered LNs, 18m cell strainer, washed, counted, and stained for flow cytometry after counterlabeling with PE conjugated anti-CD11c (Biolegend). Prior to FITC painting, some cohorts of mice were treated with simvastatin i.p. at 0.57 mg/kg/day for 7 days, as this protocol was previously shown to significantly block monocyte diapedesis from the bloodstream^51^. Control mice received vehicle i.p.

### Data availability

The bulk RNA-sequencing data are deposited in the GEO repository under the accession codes GSE122108 and GSE15907. Interactive gene expression heatmaps for both ImmGen and Tabula Muris Senis datasets as well as metabolic modules described in the present research are available by the provided link https://artyomovlab.wustl.edu/immgen-met/.

## Supplementary information

Supplementary Figures 1–9, Supplementary Tables 1–5 and Supplementary Material.

## END NOTES

## Acknowledgments

We thank Amanda Swain, Monika Bambouskova and Laura Arthur for constructive criticism of the manuscript. This work was supported in part by the Division of Intramural Research of the NIAID, NIH. The work was also partly supported by R01-AI125618 (NIAID) to M.N.A.

## Author contributions

A.S., M.N.A., A.G. designed the study and interpreted results.

A.G. performed all bioinformatic experiments and draw figures.

L.-H.H. hold the DC migration experiment.

L.S.Y., A.K., B.J, K.Kr., J.W., K.Cr., E.T., R.D., P.S., C.S., S.G., G.B., J.V.D., B.M., S.T., K.Ki., P.W., S.V.A., V.A., J.M.B., K.Ch., M.D., I.D., E.L.G., C.J., K.L., M.S.L., H.P., M.H.S., L.P., S.D., T.-A.Z. performed or participated in fluorescence-activated cell sorting as part of ImmGen MNP OS project.

C.B. and M.M. oversaw the ImmGen MNP OS logistics and overall data/sample collection.

K.S. processed raw ImmGen MNP OS RNA-seq data and produced gene counts table.

H.T. performed ImmGen MNP OS gene counts table normalization.

M.N.A., A.G., M.G., F.G., M.M., G.J.R., V.N. and A.S. participated in study discussion and provided critical insights to the study.

M.N.A. and A.G. wrote the manuscript and all the authors contributed to editing and suggestions.

## Competing interests

Authors declare no conflicts of interests.

## Notes

https://www.ncbi.nlm.nih.gov/geo/query/acc.cgi?acc=GSE122108

https://www.ncbi.nlm.nih.gov/geo/query/acc.cgi?acc=GSE15907

https://figshare.com/projects/Tabula_Muris_Senis/64982

http://artyomovlab.wustl.edu/immgen-met/

### Competing Interest Statement

The authors have declared no competing interest.

