## Supplementary Methods and Figures for "Open Source ImmGen: network perspective on metabolic diversity among mononuclear phagocytes"

##### GAM-clustering

The algorithm for multisample metabolic network clustering (hereinafter referred to as GAM-clustering) identifies modules describing dynamic regulation of metabolism and is based on the previously developed GAM method[12]. GAM-clustering extends the GAM method by setting the task to find not one but several metabolic modules (connected subnetworks of metabolic network) with the condition that each of these modules should contain as many metabolic genes with high pairwise correlation of their expression as possible.

The metabolic network used in the current analysis is presented as a graph where vertices are metabolites and edges are KEGG database reactions which are mapped with catalysing them enzymes and corresponding genes. This network is an undirected pseudograph. Totally, network contains all possible biological reactions documented in KEGG database. Reactions specific for metabolism of *Mus musculus* were selected based on gene annotation provided by KEGG and Bioconductor.

The initial approximation of the final metabolic modules is carried out by *k*-medoids clustering of the expression matrix of all metabolic genes of the dataset with some arbitrary parameter $k$ (here used $k$=32). Each cluster forms a corresponding expression pattern which can be determined as the averaged value of *z*-normalized gene expression values in this cluster. Then, a gene's $\mathrm{score}$ relative to each cluster is calculated according to formula (4). This $\mathrm{score}$ represents similarity of gene expression with the module's pattern (1) and dissimilarity with other modules' patterns (3). Formally, $\mathrm{score}$ is defined as follows:

$$d(g_i, c_j) = 1 - \mathrm{cor}(g_i, c_j) \quad (1),$$

$$d(g_i, c_0) \equiv base \quad (2),$$

$$d'(g_i, c_j) = \min_{k \neq j, k \in (0,M)} (d(g_i, c_k)) \quad (3),$$

$$\mathrm{score}(g_i, c_j) = -\log \frac{d(g_i, c_j)}{d'(g_i, c_j)} \quad (4),$$

where $g_i$ – expression of $i$-*th* gene, $i \in (1, N)$;

$c_j$ – pattern of $j$-*th* cluster, $j \in (1, M)$; $c_k$ – pattern of $j$-*th* cluster or fake pattern, $j \in (0, M)$;

$c_0$ – fake pattern;

$d$ – distance to the pattern $\mathrm{score}$ is being calculated for;

$d'$ – distance to the pattern which this gene has the most correlation with (all other patterns except the pattern the $\mathrm{score}$ being calculated for are considered);

$base$ – distance to fake pattern.

The following approach allows to avoid collapsing similar modules with enough supporting genes into one module as only one positive $\mathrm{score}$ per gene is possible.

Thus, a set of networks where each edge is weighted according to its gene $\mathrm{score}$ is formed. For each pattern a connected subgraph of maximal weight is found. These subgraphs are called metabolic modules. This procedure is carried out by a SGMWCS (signal generalized maximum weight connected subgraph) solver[19,49] (https://github.com/ctlab/sgmwcs-solver) which uses the IBM ILOG CPLEX library that efficiently performs many iterations of this method in a reasonable amount of time. Thus, an iterative procedure of metabolic modules refinement is performed in a process of updating each of the patterns by replacing it with an averaged gene expression of the module's edges with a positive $\mathrm{score}$.

One of the important parts in the procedure of updating the modules is the question when to stop. To detect this, the difference between the values of the patterns of the current iteration and the values of the patterns of all previous iterations, in which there were the same number of modules, is found (this is done to avoid missing the situation when new iteration comes to the condition close to one that once already has occurred). If difference is large (>0.01) which means that pattern content is quite changed, a new score set is calculated and a new iteration is performed. If the difference between patterns is small enough (<0.01), but non-informative (having less than 5 edges and/or diameter less than 4) modules are still presented in the output, the less informative (most correlated with any other graph) module is eliminated from the further analysis. After removing one module, the weights are recalculated and a new iteration of refinement is performed. The final result is a set of specific subnetworks that reflect metabolic variability among the samples of the analysed transcriptome data.

The GAM clustering method has two parameters: the number of initial clusters $k$ (here used $k$=32) and the distance to the fake pattern $base$ (here used $base$=0.4). They directly affect the number, size, intramodular characteristics, and the number of unique annotating pathways of the resulting modules (**Supplementary Fig. 5a,b**).

To explore the influence of $k$ value to number of final modules the model data were designed. They imitate experiment with complex design (15, 18 or 21 samples) where several (5, 10 or 15) modules are active each in a particular subset of samples. All combinations of these data were analysed by the GAM-clustering method and the following output features were calculated: number of final modules found by method, number of iteration performed and time elapsed during the analysis (**Supplementary Fig. 5a**). As these data were modelled, we know how many modules are there in each experiment (dashed line in **Supplementary Fig. 5a**) and therefore we can evaluate how the number of found modules relates to the number of real modules. In most cases GAM-clustering found approximately all real modules when launched with the value $k$ several times greater that the number of real modules. Moreover, a further increase of $k$ does not lead to improved results, but nonlinearly increases the number of iterations and the working time of the method. Thus, it is reasonable to detect some advisable $k$ value so that user gets approximately full set of modules and does not spend to much time for the analysis. As in real data we do not know the number of real modules there is a heuristic approach that allows to find some $k$ based on the characteristics of the input data. This approach is based on elbow method that calculates the total within-cluster sum of square (wss) for each $k$. As expression data have significant noise contribution there is no pronounced inflection point where wss is sharply stops decreasing (usually this point is considered equal to the optimal number of clusters). Here, we used point where the slope of the wss curve is 50% of its steepest slope. Corresponding to this point value of $k$ is rounded to the nearest value used in the practice (16, 24, 32, 40, 48, 56, 64), and the obtained value is proposed by the method as the recommended $k$ value.

The strategy for selecting optimal value of $base$ parameter was formed on the basis of real data analysis, since it requires consideration of the biological meaning of the obtained modules. At the beginning of the analysis, the GAM-clustering algorithm produces some recommended value of $k$ (see previous paragraph). For this $k$, we can calculate the average dissimilarity (distance) between the observations of the initial cluster and this cluster's medoid over all clusters. Obtained value is proposed by the method as the recommended value of the $base$ parameter. For the ImmGen MNP OS data analysed in this study, they were 32 initial clusters proposed and the recommended $base$ value was equal to 0.4. This $base$ value was determined to be optimal during the comparative study of the results obtained with other different $base$ values (**Supplementary Fig. 5b**). The optimality criterion included the calculation of the following characteristics of the output modules: their number, size, average correlation of edges, the number of unique annotating paths, the number of annotating paths corresponding to one cluster only, the percentage of genes with negative $\mathrm{score}$, the percentage of genes with negative correlation, the percentage of genes with correlation less than 1-$base$. Noticeably, such characteristics as the average number of genes in the module, the average percentage of genes with negative $\mathrm{score}$ and correlation, as well as with a correlation less than $1 - base$, are minimal for the recommended $base$ value (0.4). This indicates that the modules obtained for $base$ = 0.4 have good internal correlation, as well as compactness. Modules obtained with a lower $base$ value also show good internal correlation, but they are characterized by the loss of a large number of significant modules. It is worth noting that for $base$ = 0.2 no modules were found. Modules obtained with larger $base$ values, on the contrary, are annotated with a bigger number of unique canonical pathways, however, many of these pathways relate to the same biochemical processes. Moreover, these modules are characterized by lower rates of intramodular correlation.

Even though default values of $k$ and $base$ parameters are proposed to user before the analysis based on the input data properties, there is still an opportunity for user to select custom values of these parameters. Nevertheless, the general recommendation is to stick with the proposed value of the $base$ parameter, since its changes lead to the strong alterations in the size and content of the final modules.

GAM-clustering method is applicable not to bulk RNA-seq data only but to single-cell RNA-seq data as well. Single-cell data need an additional step of preprocessing implying transformation of individual cells into technical samples. This is performed based on averaging gene expression of individual cells inside high resolution clusters. In case of single-cell RNAseq data, among final metabolic modules might occur ones that do not cover all biological replicas of cell types they are specific for. These modules are eliminated from the final result.

The final metabolic modules are subnetworks of the overall metabolic network that contain a set of closely located genes with high correlation of their expression profile across all samples.

Code is available at https://github.com/artyomovlab/ImmGenOpenSource.

**SUPPLEMENTARY FIGURE LEGENDS**

**Supplementary figure 1.** Principal component analysis (PCA) of ImmGen MNP OS dataset based on 12,000 most expressed genes across all samples colored by lab of samples sorting **(a)** and batch of samples sequencing **(b)**. Axes are the first two principal components (PCs). **c,** Boxplots of either raw either normalized counts of 12,000 most expressed genes across all samples.

**Supplementary figure 2.** Principal component analysis (PCA) of ImmGen MNP OS dataset based on 12,000 most expressed genes across all samples colored by cell specific markers expression (from lowest as blue to highest as red). Axes are the first two principal components (PCs).

**Supplementary figure 3. a,** Principal component analysis (PCA) based on 12,000 most expressed genes across all samples colored by intensity of gene expression (from lowest as blue to highest as red) of several KEGG and Reactome canonical pathways for ImmGen MNP OS **(a)** and ImmGen MNP P1 **(b)** datasets.

**Supplementary figure 4.** Violin plot for the number of genes in the cells of each natural cluster of mTMS dataset.

**Supplementary figure 5. Analysis of $k$ and $base$ parameter values influence to characteristics of final modules performed on model (a) and ImmGen MNP OS (b) data. a,** Model data imitate experiment with complex design (15, 18 or 21 samples; shown by row splitting) where several modules (5, 10 or 15; shown by colored dashed line in the first column) are active each in a particular subset of samples. All combinations of these data were analysed by the GAM-clustering method with $k$ values equal to 16, 24, 32, 40, 48, 56 or 64 and the following output features were calculated: number of final modules found by method (first column), number of iterations performed (second column) and time elapsed during the analysis (third column). **b,** Comparative study of the results obtained after GAM-clustering analysis of the ImmGen MNP OS data (with 32 initial clusters) with different $base$ values (0.3, 0.4, 0.5 or 0.6). Optimal value for the $base$ parameter (framed) was determined by the calculation of various characteristics of the output modules.

**Supplementary figure 6.** Principal component analysis (PCA) of ImmGen MNP OS dataset in module **(a)** and transcriptional **(b)** spaces across all samples colored on the basis of its belonging to a particular metasample (metasample names are given based on the major cell type in the current metasample: Mo – monocyte, MF – macrophage, DC – dendritic cell, YS MF – yolk sac macrophage, EB MF – embryoid body macrophage, alvMF – alveolar macrophage, SPM – small peritoneal macrophage, MG – microglia, pDC – plasmacytoid dendritic cell, migDC – migratory dendritic cell).

**Supplementary figure 7.** UMAP plots colored by intensity of gene expression (from lowest as blue to highest as red) of modules derived from GAM-clustering analysis of mTMS dataset; prog – progenitor, MF – macrophage, alvMF – alveolar macrophage, MG – microglia, KC – Kupffer cell, Mo – monocyte, DC – dendritic cell, NP – neutrophil.

**Supplementary figure 8. Metabolic modules 1 (a) and 2 (b) *per se* (subnetworks associated with lipid metabolism).** Edges of modules are attributed with color according to correlation of its enzyme's gene expression to this particular module pattern and with thickness according to its score.

**Supplementary figure 9. Subnetworks associated with fatty acid synthesis in cytosol (b) and mitochondria (c). a,** Heatmaps of module patterns along with the expression of some of its genes (from lowest as blue to highest as red). **b,c,** Metabolic modules *per se* and corresponding schematic diagrams. Edges of modules are attributed with color according to correlation of its enzyme’s gene expression to this particular module pattern and with thickness according to its score. **d,** Enrichment of modules genes expression (from lowest as blue to highest as red, transparent dots correspond to treated samples) across all three analysed datasets.

### Supplementary figure 1

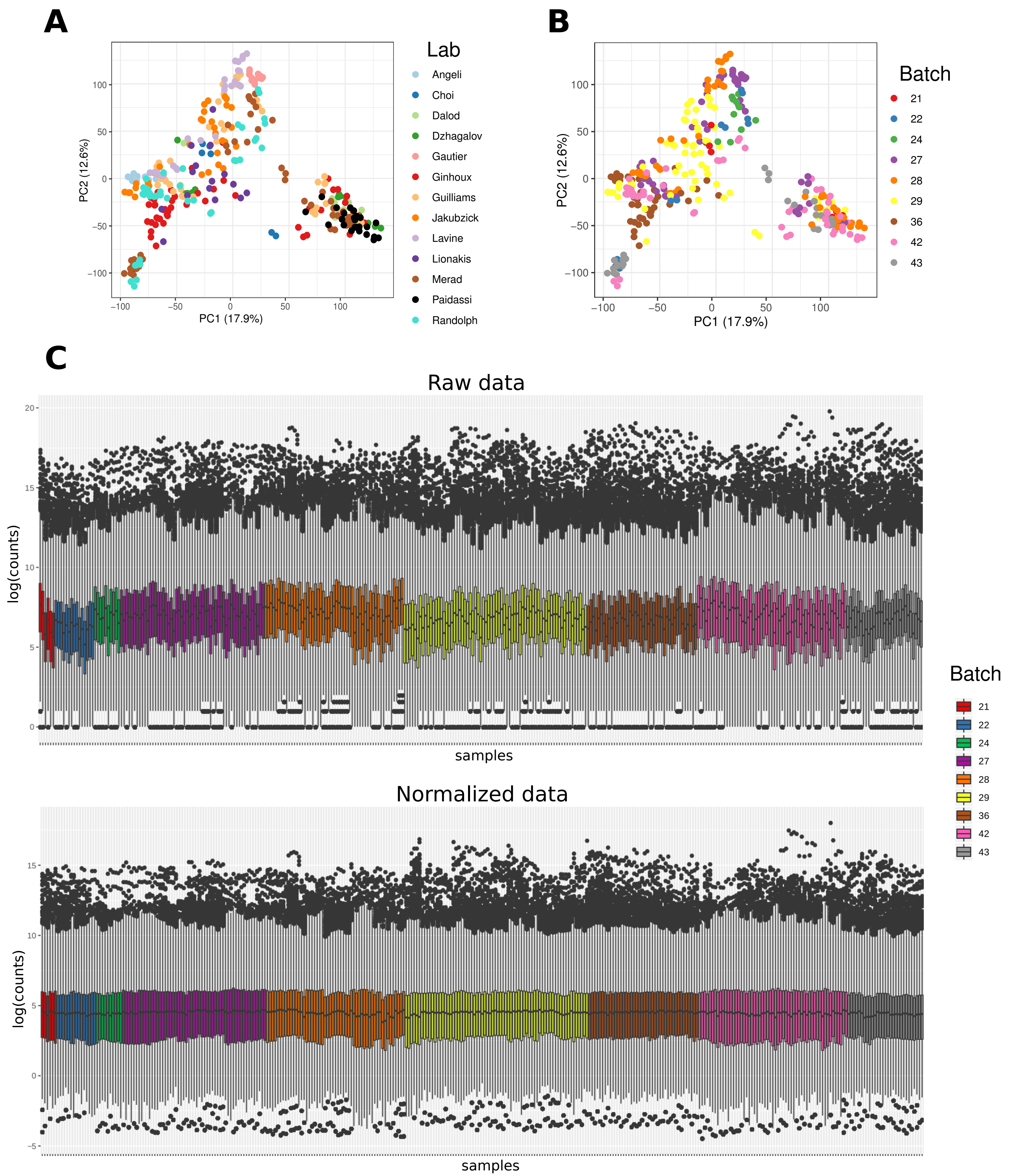

### Supplementary figure 2

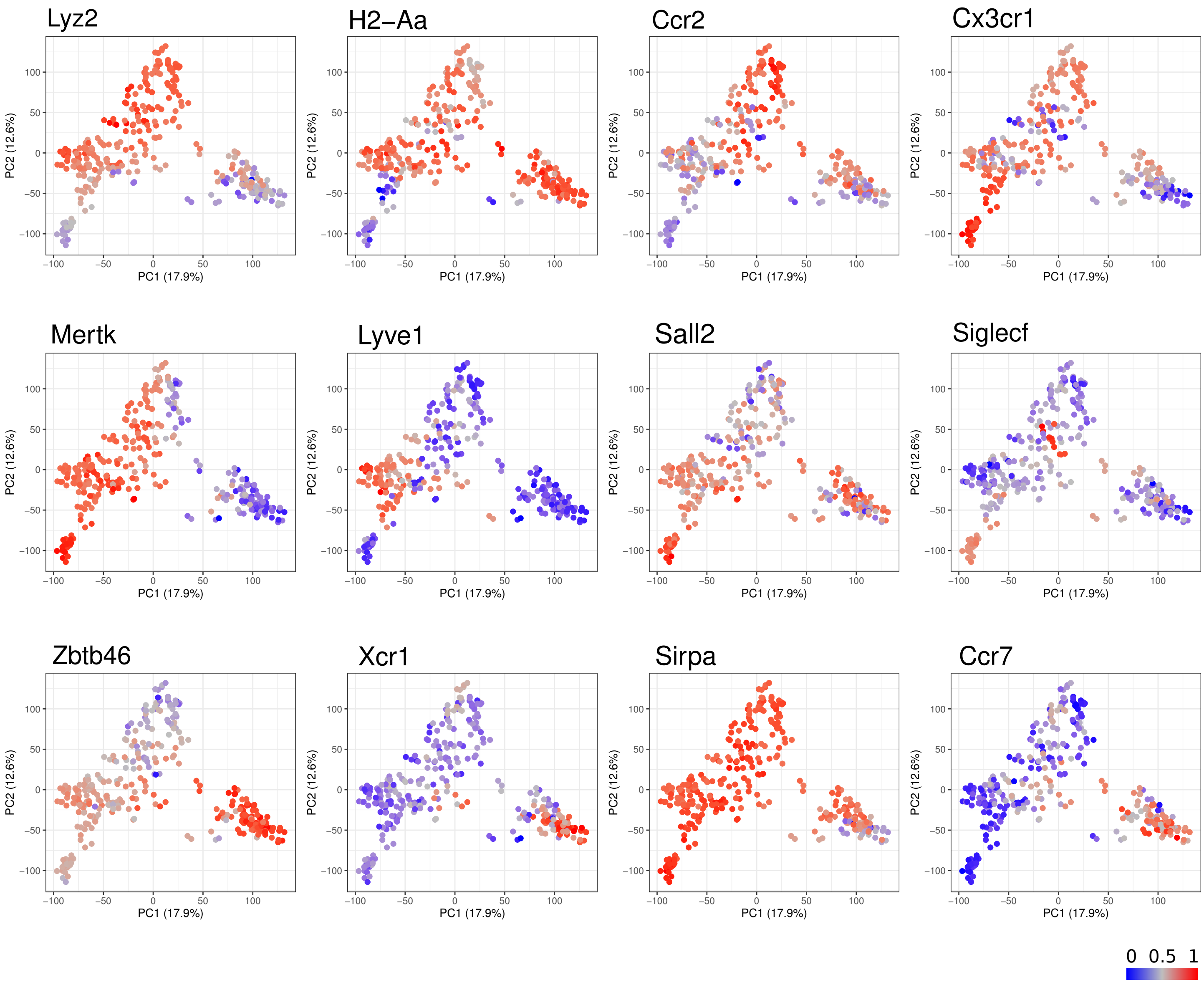

### Supplementary figure 3

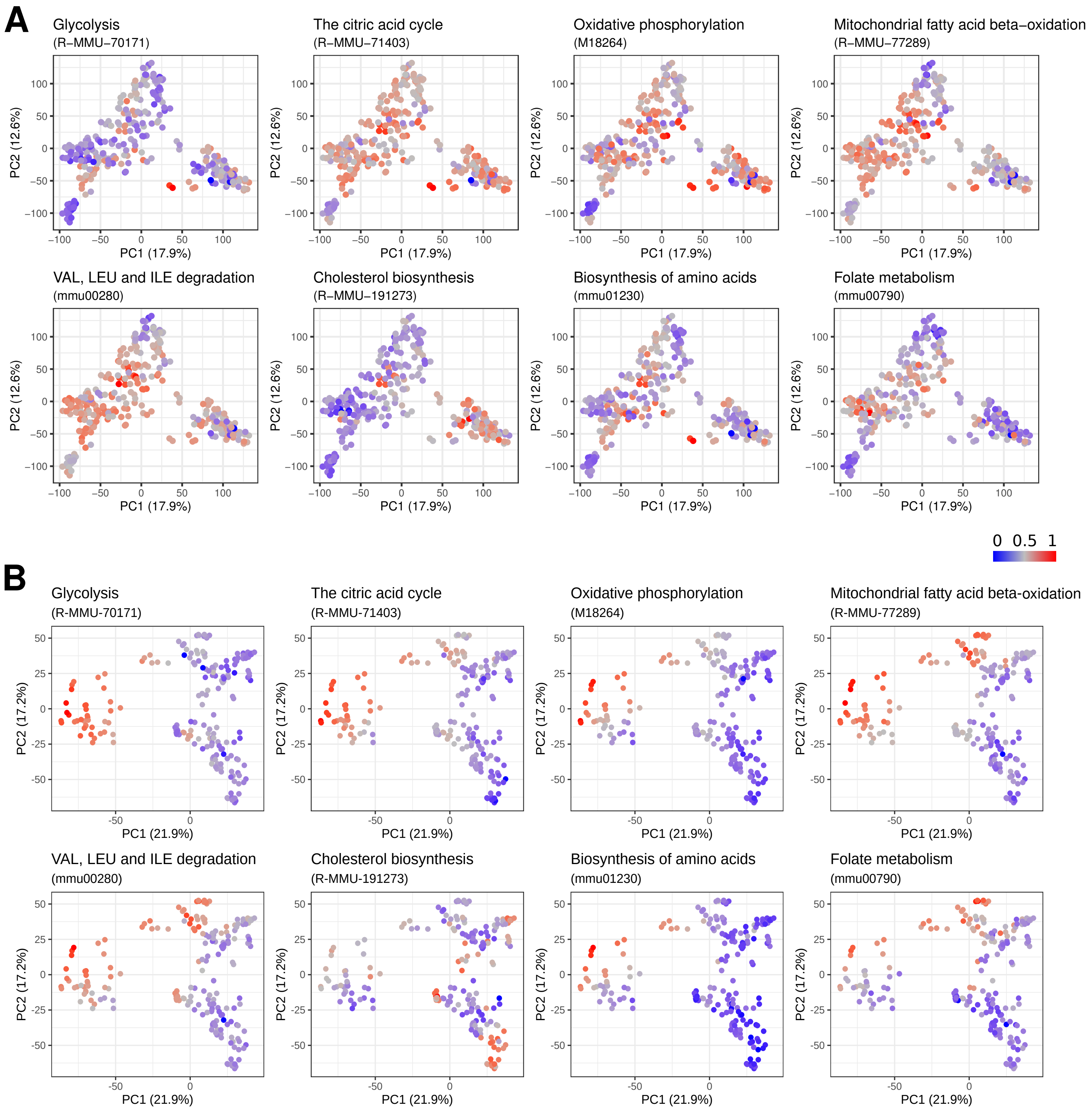

### Supplementary figure 4

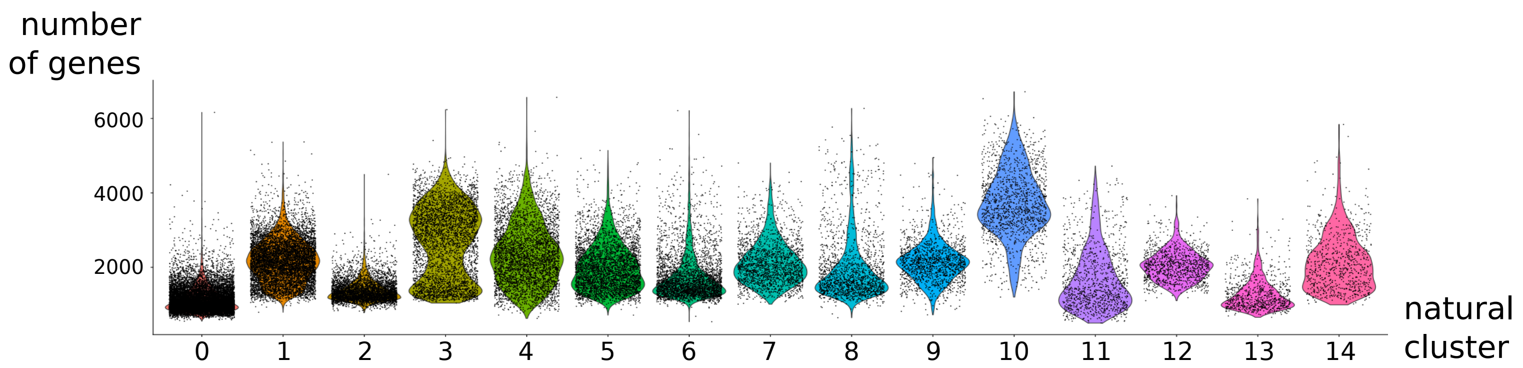

### Supplementary figure 5

**A**

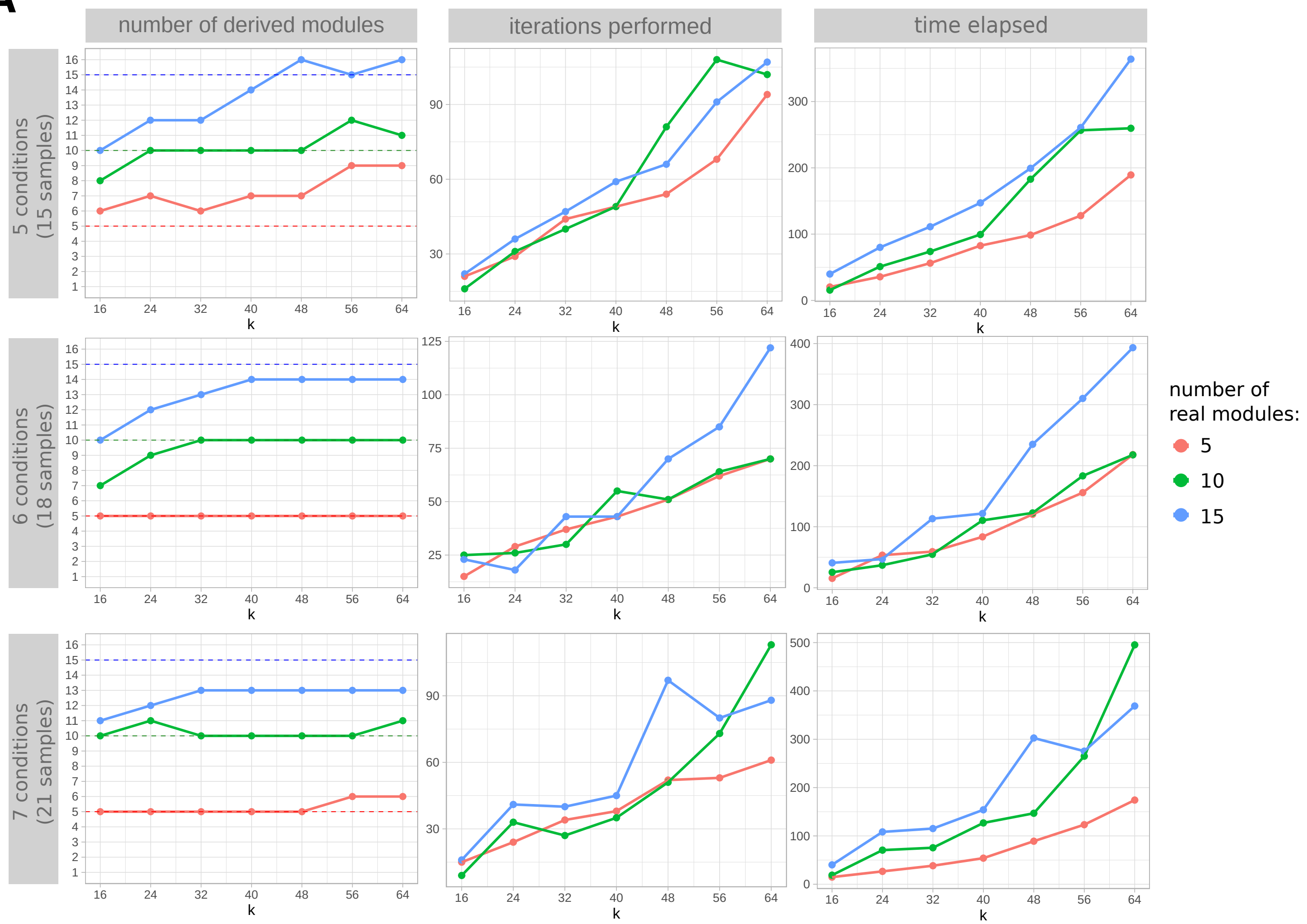


**B**

| *base* | 0.3 | 0.4 | 0.5 | 0.6 |
|---|---|---|---|---|
| number of modules | 3 | 9 | 10 | 16 |
| mean number of module edges | 24.3 | 20.9 | 33.9 | 24.6 |
| mean correlation of module edges | 0.79 | 0.74 | 0.67 | 0.64 |
| number of unique annotating pathways | 8 | 25 | 36 | 36 |
| number of pathways related to one module only | 8 | 17 | 25 | 21 |
| mean percent of genes with negative score in module | 17.4% | 12.5% | 18.7% | 19.2% |
| mean percent of genes with negative correlation in module | 0% | 0% | 0.4% | 1% |
| mean percent of genes with correlation less than 1-*base* in module | 17.4% | 9.4% | 15.5% | 12.1% |

### Supplementary figure 6

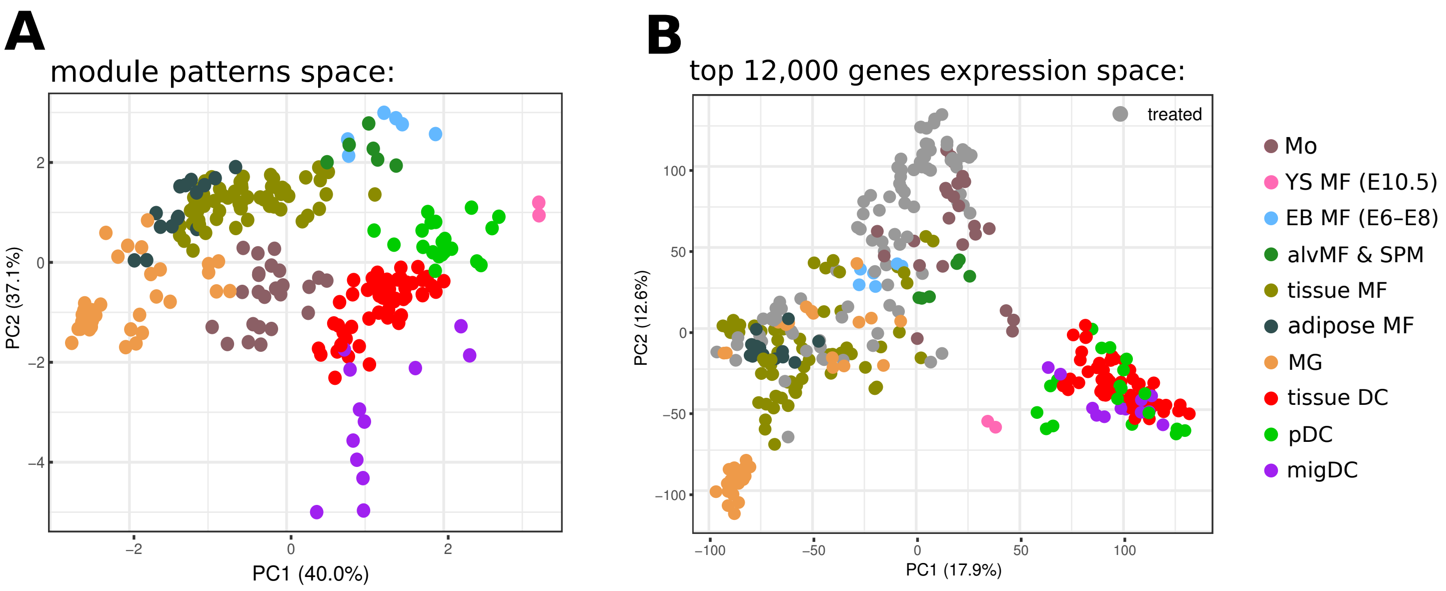

### Supplementary figure 7

modules derived from GAM-clustering analysis of mTMS dataset

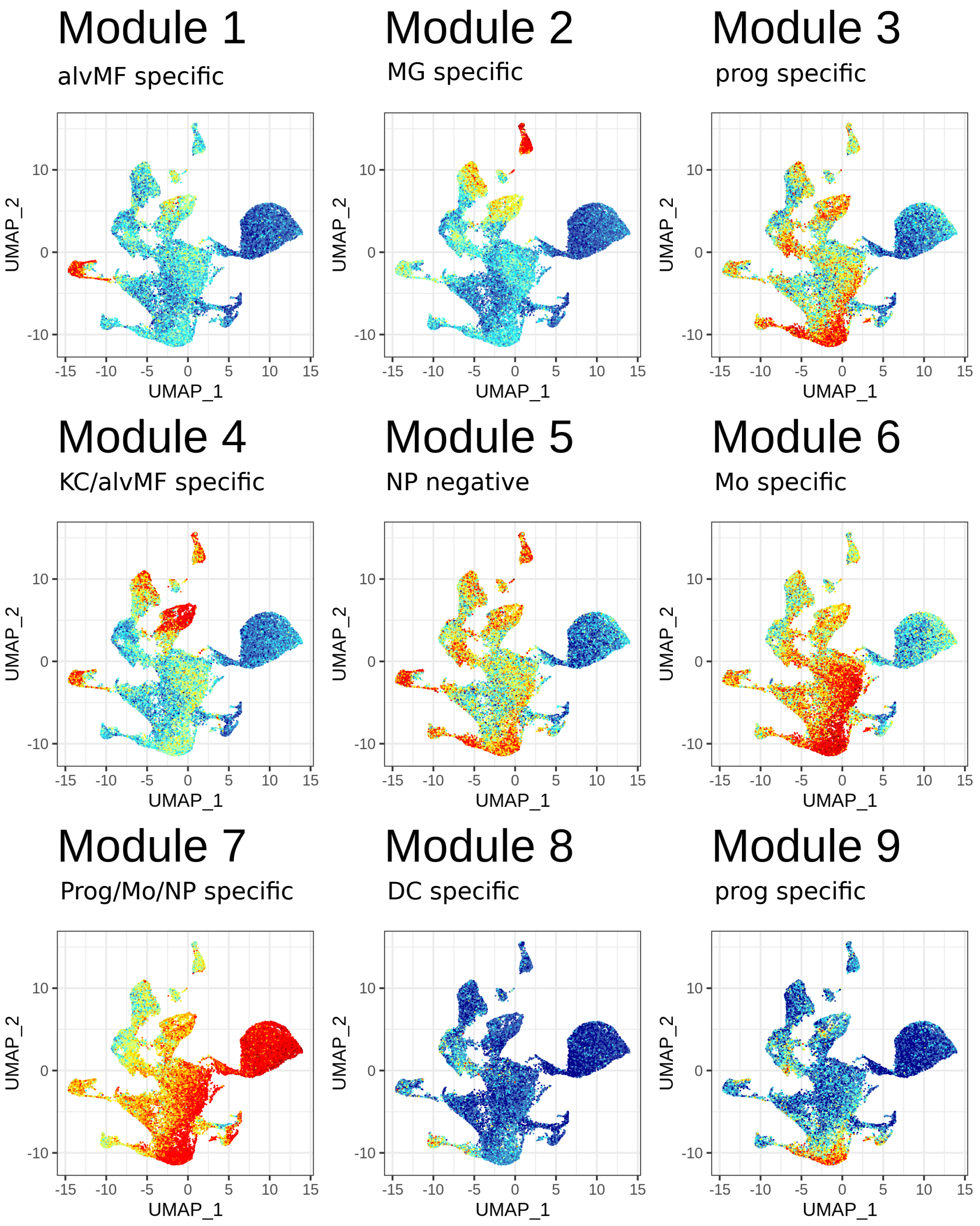

### Supplementary figure 8

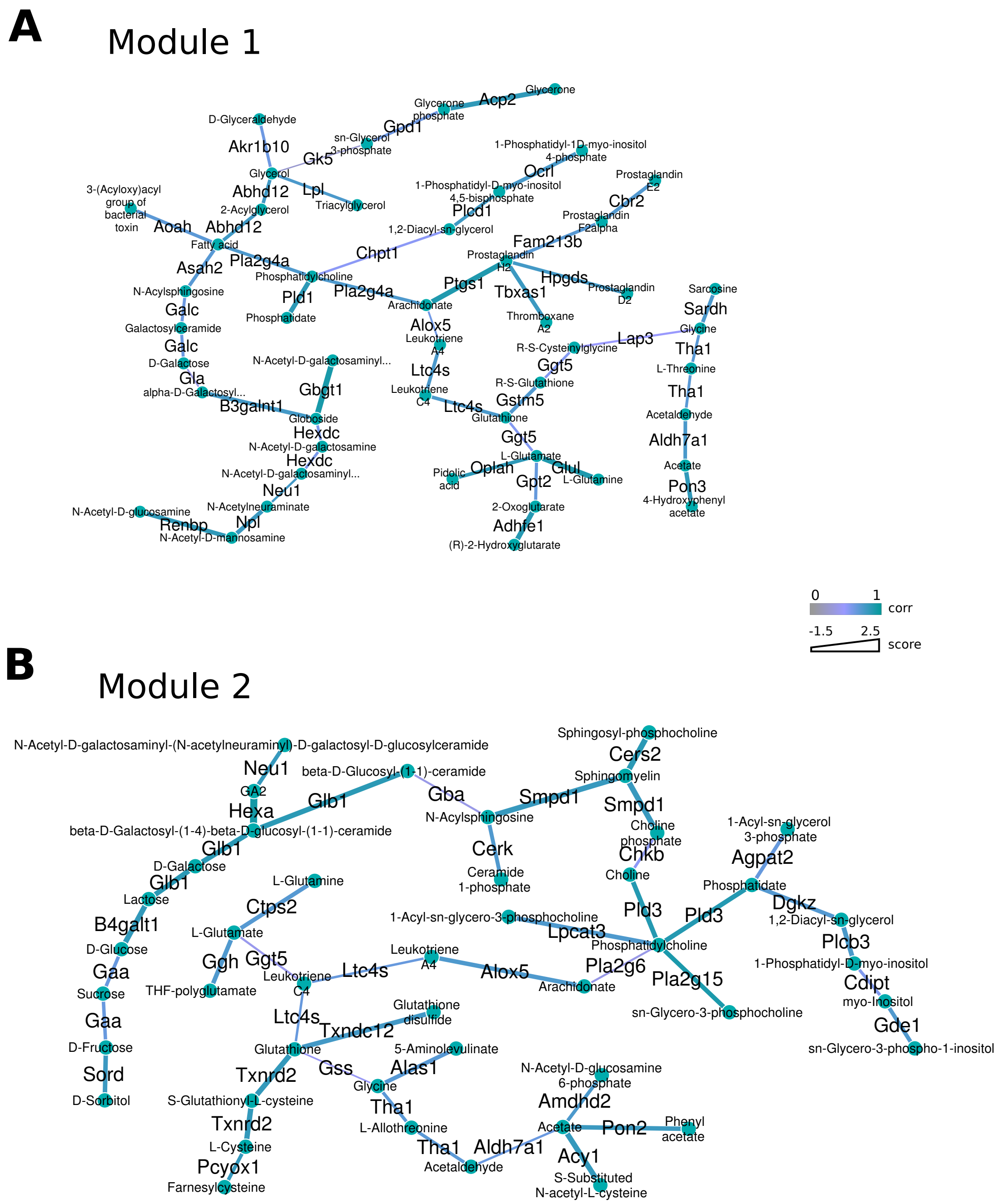

### Supplementary figure 9

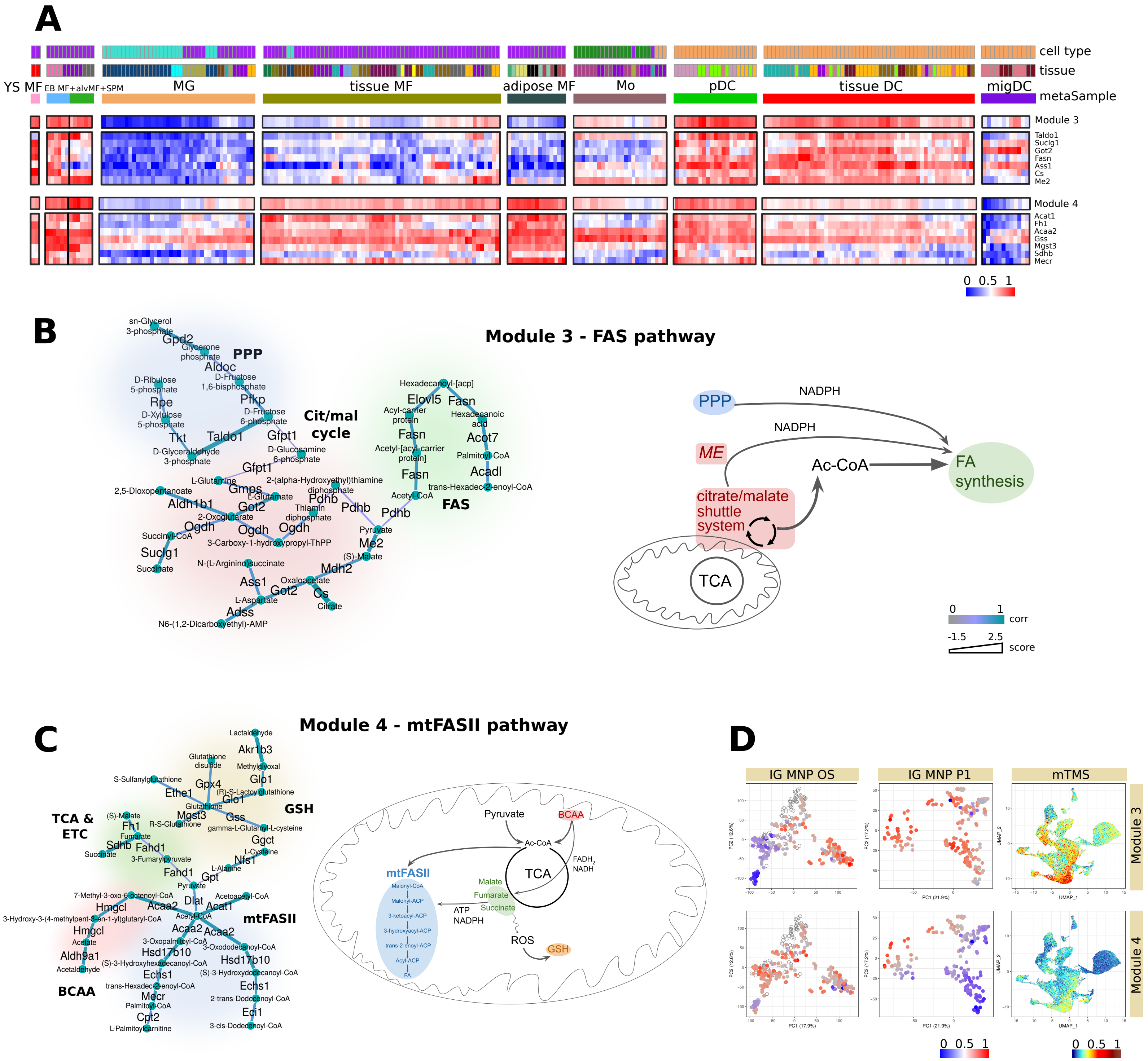
